# Image-based transposon screening reveals a flavin reductase that restrains intracellular *Salmonella* replication in macrophages

**DOI:** 10.64898/2026.07.30.741741

**Authors:** Camilla Ciolli Mattioli, Daniel Schraivogel, Noa Bossel Ben-Moshe, Hadar Ben-Arosh, Agzamara Gonzales Acosta, Suzie Rousselle Kenmoe, Sharon Ben-Hur, Lars Michael Steinmetz, Roi Avraham

## Abstract

Intracellular bacterial pathogens can survive and replicate within host cells, yet an isogenic population follows divergent fates: some bacteria are killed, some arrest growth, and others replicate to high numbers. Identifying the bacterial functions behind each fate requires recovering mutants from within the host cells displaying it. But systematic approaches, such as transposon insertion analysis, can only estimate fitness of mutants from the whole infected population. Here, we developed an approach that couples a genome-wide mutagenesis library to image-enabled cell sorting (ICS), sorting infected cells by the number of bacteria they contain and assigning mutants to defined replication outcomes. We applied this approach to *Salmonella enterica* serovar *Typhimurium* (*S.*Tm) in macrophages, and revealed genes required for replication from genes that restrain it, whose disruption increased replication. Among the latter we identified the cytosolic flavin reductase Fre, which supplies reduced flavins to a broad range of bacterial processes. We uncovered a mechanism whereby loss of *fre* protected *S.*Tm from oxidative and nitrosative damage and increased bacterial numbers. Inside macrophages this advantage was mediated by the upregulation of the iron-sulfur-independent cytochrome bd-I oxidase. By resolving a mutant library into phenotypically defined subpopulations, this framework can be applied to characterize bacterial or host genes that drive infection phenotypes in any infection model.

## Introduction

Intracellular bacterial pathogens can survive and proliferate inside host cells. Yet an isogenic population does not replicate uniformly: some bacteria replicate within a remodeled, nutrient-poor compartment [1,2], while others escape to the cytosol [3,4]. Individual bacteria also follow distinct fates within these niches, where some are killed, some arrest in a non-replicating or slowly replicating state, and others replicate to high numbers within the same host-cell population [5]. Thus, the intracellular lifestyle of bacteria is characterized by pronounced single-cell heterogeneity, which has been studied in depth in *Salmonella enterica* serovar Typhimurium (*S.*Tm) within the hostile environment of macrophages. To restrain intracellular *S.*Tm, macrophages deploy an arsenal of antimicrobial effectors, including reactive oxygen species, reactive nitrogen species, phagosomal acidification, and restriction of essential nutrients [6]. *S.*Tm counters these assaults with dedicated detoxification and repair systems [7]. However, a systematic approach to resolve which bacterial functions drive each of these distinct intracellular phenotypes of survival and replication is still lacking.

The machinery of the intracellular lifestyle has instead been mapped almost entirely by transposon insertion sequencing, which has identified genes required by *S.*Tm to survive within macrophages and animal hosts [8–22]. In these screens, a pooled mutant library is used to infect the host, and the abundance of each mutant in the recovered population is compared to the input library before infection. While these screens provide invaluable understanding of the competitive fitness of bacterial mutants during intracellular infection [8–22], they average over different infection phenotypes coexisting in the infected population. A mutant lost from the pool may have been killed, blocked from replicating, or restricted by the host, and a mutant enriched in the pool may survive better rather than replicate more. Resolving these phenotypes requires probing infection at the level of single cells, and fluorescent reporters have been central to this effort. Division-tracking reporters such as fluorescence dilution [23,24] and maturation-based TIMER [25] systems distinguish replicating from non-replicating bacteria; transcriptional fluorescent-protein reporters report several types of stress responses [26,27], as well as cytoplasmic or vacuolar localization [28]. Together these tools have established that an isogenic *S.*Tm population diversifies into subpopulations with distinct growth rates, stress exposure, and fates. These reporters, however, are usually read out by flow cytometry on bacteria liberated from their host cells, so that each event is a single bacterium detached from its macrophage, and both the number of bacteria per infected cell and any host-associated phenotype are lost [25,27]. Infected cells can instead be analyzed intact, by gating on macrophages and normalizing the reporters against a second fluorophore [24,29], an approach also coupled to single-cell RNA-sequencing of the host to link macrophage polarization state to bacterial growth rate, yet the integrated signal is continuous, and thus bacterial number cannot be resolved into discrete per-bacterium steps from intensity alone [30,31]. Imaging intact infected cells overcomes both limitations: it preserves the host cell as the unit of analysis and resolves individual bacteria. Imaging further captures information beyond bacterial number: the arrangement of intracellular bacteria and host-derived features such as cell morphology [32] and the subcellular localization of the infection [28,33], which report on phenotypes ranging from bacterial replication and localization [33] to engagement of host cell death [34,35]. Yet single-cell approaches such as these resolve bacterial fate without revealing its genetic basis, just as transposon screens assign genes to fitness while averaging across those same fates. What is now needed are approaches that combine mutant screens with imaging of selected phenotype of infection, for example, by determining the non-replicating and replicating subpopulations that coexist within the same infection.

Image-enabled cell sorting (ICS) [36] brings image-level features into a sortable readout, combining per-cell imaging with high-speed sorting so that infected cells can be characterized and physically recovered on the basis of image-derived features. Here, we developed an approach, based on ICS [36], that couples single-cell imaging and sorting to a genome-wide mutagenesis library. We applied this approach to dissect genes that are required for the phenotype of intracellular replication of *S.*Tm. We infected macrophages with a mutant library of *S*.Tm, and using ICS sorted infected cells based on the number of bacteria they contain. We optimized protocols for intracellular recovery of bacterial transposon identity and recovered mutants required for replication and mutants that restrain it, whose disruption increased the number of bacteria per macrophage. Through this analysis we identified novel genes and pathways involved in intracellular infection, including the cytosolic flavin reductase Fre as a restraint on intracellular replication. Rather than protecting *S.*Tm, Fre potentiated the oxidative and nitrosative damage, so that its loss increased bacterial replication. We suggest that Fre exemplifies a bacterial function that is counterproductive within the macrophage environment, linking flavin metabolism to the redox chemistry of intracellular survival.

## Results

### Image-enabled cell sorting separates macrophages by intracellular bacterial load

Macrophages that internalize similar numbers of bacteria nonetheless diverge over the course of infection, some containing a single non-dividing bacterium and others actively dividing cells [24,37]. To dissect the bacterial determinants of replication heterogeneity, we first needed to separate infected macrophages by the number of bacteria they contain. To do so, we established a strategy based on high-speed image-enabled cell sorting (ICS) [36], which makes high-speed measurements on multicolor fluorescence images, allowing cells to be sorted based on these image-derived measurements. We applied ICS to bone marrow-derived macrophages (BMDMs) infected with *S.*Tm at multiplicity of infection (MOI) of 2:1. Intracellular bacteria were visualized by anti-LPS O-antigen immunostaining (FITC). We defined a gating strategy within infected cell populations using image-derived measurements and traditional flow cytometry parameters (**Fig. 1a** and **S1a,b**). For the low-load gate (1 bacterium per cell), cells infected with a single bacterium produced a compact, centrally located FITC spot of modest brightness. We were able to isolate these cells using the imaging parameter FITC diffusivity and FITC-A. For the high-load gate (≥5 bacteria per cell), bacteria occupy a larger spatial footprint within the host cell, which we resolved using the imaging parameters FITC size and FITC max intensity.

**Fig. 1:**
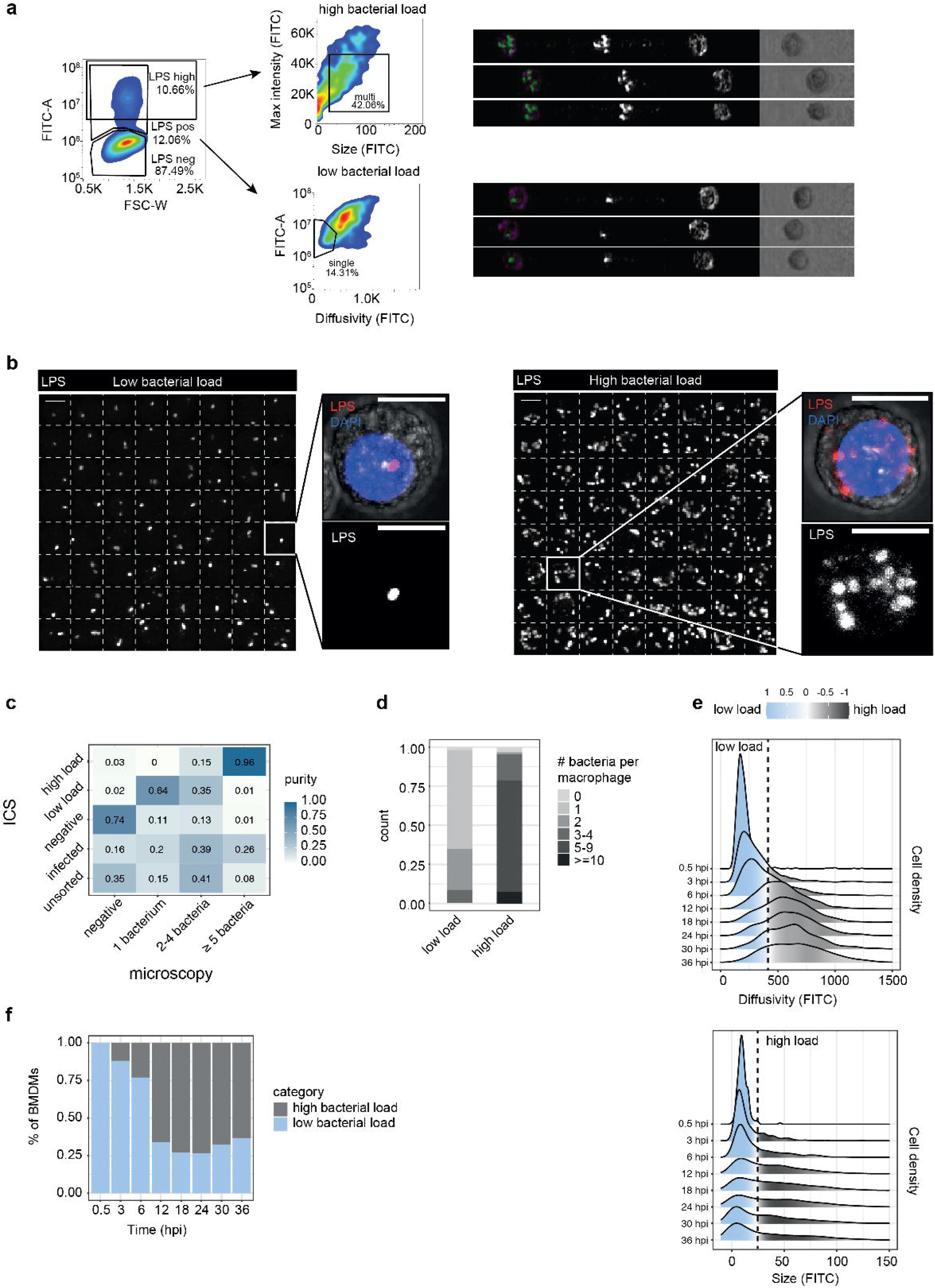
Image-enabled cell sorting separates infected macrophages by intracellular bacterial load. **a,** Image-enabled cell sorting (ICS) separates infected macrophages by intracellular bacterial load. Gating strategy for BMDMs infected with anti-LPS–stained (FITC) *S.*Tm. LPS^+^ infected cells were split into a high-load gate, defined by FITC max intensity versus FITC size, and a low-load gate, defined by FITC-A versus FITC diffusivity. Right, representative ICS images of high-load (top) and low-load (bottom) events. **b,** Sorted populations match their assigned bacterial load by traditional confocal fluorescence microscopy. Montages of low-load (left, 1 bacterium per cell) and high-load (right, ≥5 bacteria per cell) sorted cells (LPS channel), with magnified insets showing a representative single-bacterium and multi-bacteria macrophage. LPS, red; DAPI, blue. Single channels are shown in black. Scale bar, 10 μm. **c,** ICS gates are pure for their intended bacterial load. Classification matrix comparing ICS gate assignment (rows) with bacterial load scored by microscopy (columns). Each row is normalized to the total events in that population; values and color give the fraction assigned to each microscopy category. 131-357 cells were scored per population. The “infected” and “unsorted” rows show the composition of the bulk-infected and pre-sort populations as a reference baseline. **d,** The low- and high-load gates are enriched for increasing numbers of bacteria. Distribution of bacteria per macrophage, scored by microscopy, within each ICS gate. **e,** Intracellular bacterial load shifts from low to high over the course of infection. Density distributions of FITC diffusivity (top) and FITC size (bottom) for infected macrophages, measured by ICS at 0.5–36 hpi (MOI 2:1). Ridgeline height shows the overall cell density at each position; fill color reflects, among the cells at that position, their membership in the two-parameter low- and high-load gates projected onto the displayed axis, computed as the difference between the kernel-weighted proportions of low-load and high-load cells: light blue, predominantly low-load; grey, predominantly high-load; near-white, cells not strongly assigned to either gate. The dashed line marks the projected gate boundary. Each time point is indicated on the left. **f,** The infected population remains heterogeneous throughout infection. Proportion of BMDMs classified as high- versus low-bacterial-load by ICS at 0.5–36 hpi.

To benchmark the purity of these subpopulations, we infected BMDMs with *S*.Tm at MOI 10:1 to generate a broad distribution of intracellular loads, sorted the two populations based on the above gating strategy, and analyzed sorted cells by microscopy (**Fig. 1b**). The high-load gate reached 96% purity with a median of 6 bacteria per cell; the low-load gate, selected from multiple candidate imaging parameters as the one with the highest purity (**Fig. S1c**), achieved 64% purity with a median of 1 bacterium per cell (**Fig. 1c,d** and **S1c,d**). The residual impurity was almost entirely accounted for by events containing exactly two bacteria (**Fig. 1d**), the expected consequence of the axial (z) resolution limit of image-enabled cytometry, in which two bacteria stacked along the imaging axis project onto a single spot and cannot be resolved without optical sectioning (**Fig. S1e**). This may reflect co-occupancy of a single vacuole by two bacteria, a transient phenomenon documented during *S.*Tm infection [38,39].

To confirm that infection at low MOI yields predominantly single-bacterium entries, we infected BMDMs at MOI 2:1 with a pool of six *S.*Tm strains, each expressing a spectrally distinct fluorescent protein, and quantified the number of colors per infected macrophage as a proxy for the number of independent bacterial entries. Only ∼5% of infected macrophages harbored bacteria of more than one color at 30 minutes post-infection (**Fig. S1f**), confirming that the majority of cells were initially infected by a single bacterium. We then tracked bacterial loads over 36 hours by ICS at MOI 2:1 to favor single-bacterium infections (**Fig. 1e,f** and **S1g**). At 0.5 hpi macrophages fell exclusively within the low-load gate. Replication initiated by 3 hpi and plateaued around 18 hpi, with the population remaining heterogeneous at all subsequent time points, comprising both low- and high-burden cells.

Together, these results established ICS as a quantitative readout of intracellular bacterial load and revealed sustained heterogeneity in *S.*Tm intracellular replication, with a subpopulation of bacteria establishing a replicative niche and another remaining as a single non-replicating infecting bacterium.

### ICS-coupled transposon sequencing identifies bacterial determinants of intracellular replication

We next sought to identify the bacterial determinants underlying the replication heterogeneity revealed by ICS. We infected BMDMs with a saturated *S.*Tm transposon mutant library at MOI 2:1. We analyzed infected macrophages by ICS, sorted into low- and high-load populations at 20–24 hpi, and processed for Transposon Directed Insertion Site Sequencing (TraDIS).

To analyze mutants enriched within each population (non-replicative and hyper-replicating), we sought to optimize bacterial gDNA library preparation directly from sorted macrophages. Current protocols for mutant identification within infected cells entail harvesting intracellular bacteria and re-growing them in liquid culture to obtain sufficient bacterial gDNA. However, regrowth introduces lag-time artifacts: bacteria recovering from macrophage stress resume growth at variable rates [24,40], and small lag differences are exponentially amplified into large differences in mutant abundance. Direct extraction circumvents this problem but yields mixed host (murine) and bacterial gDNA, which initially limited our transposon read recovery. We therefore redesigned the adapter architecture and added a nested PCR primed within the transposon to selectively amplify genuine transposon-genome junctions, a protocol we termed nested TraDIS (nTraDIS) (**Fig. S2a**). Technical replicates of the nTraDIS library were highly reproducible (R=0.98, compared to R=0.69 for the TraDIS library, **Fig. S2b**). nTraDIS also increased on-target read recovery from ∼10% to ∼85%, with mouse-aligned reads dropping from ∼64% to ∼1% (**Fig. S2c**), and recovered ∼4-fold more unique transposon insertions (**Fig. S2d**, left panel), while reducing spurious library products from ∼3.5% to ∼0.3% (**Fig. S2d**, right panel). We further characterized the starter inoculum library to confirm its quality. Genes previously annotated as essential [41] were largely devoid of insertions (**Fig. S2e**). The few insertions they carried fell toward the 3′ end of the coding sequence (**Fig. S2f**, left panel), where they are least likely to abolish function, whereas non-essential genes were covered uniformly (**Fig. S2f**, right panel). With ∼57 insertions per kilobase of coding sequence, these features confirm a saturated library suitable for genome-wide fitness scoring.

Applying nTraDIS to the ICS-sorted macrophage populations identified mutants differentially enriched between the non-replicative and hyper-replicating populations (**Fig. 2a**). In total, we identified 400 positive and 45 negative determinants of replication, enriched in the non-replicative and hyper-replicative populations, respectively (Supplementary Table S1). Pathway and regulon enrichment analysis recovered known intracellular fitness determinants (**Fig. 2b**). Mutants in *Salmonella* pathogenicity islands 1 and 2 (SPI-1, SPI-2) were strongly enriched in the low-load gate. SPI-2 has a well-established role in intracellular replication [42–44]. SPI-1, although traditionally linked to epithelial invasion [45,46], is also recognized to act inside macrophages: SipA targets host mitochondrial dynamics [47], and T3SS assembly itself dampens host immune responses through translational reprogramming [48]. Auxotrophies in essential metabolite biosynthesis were also strongly represented, including histidine (*hisA-Q*), purine (*purD*, *purL*), and mannose (*manA*) pathways, consistent with prior work establishing these as macrophage-replication determinants [49,50]. Mutants in O-antigen and LPS biosynthesis (*rfa*, *rfb*, *waa* loci) were enriched in the non-replicative population, consistent with their established roles in intracellular fitness [10,51]. However, as the efficiency of the anti-LPS O-antigen antibody used for staining depends on intact O-antigen, their low-load classification cannot be unambiguously attributed to an intracellular replication defect.

**Fig. 2:**
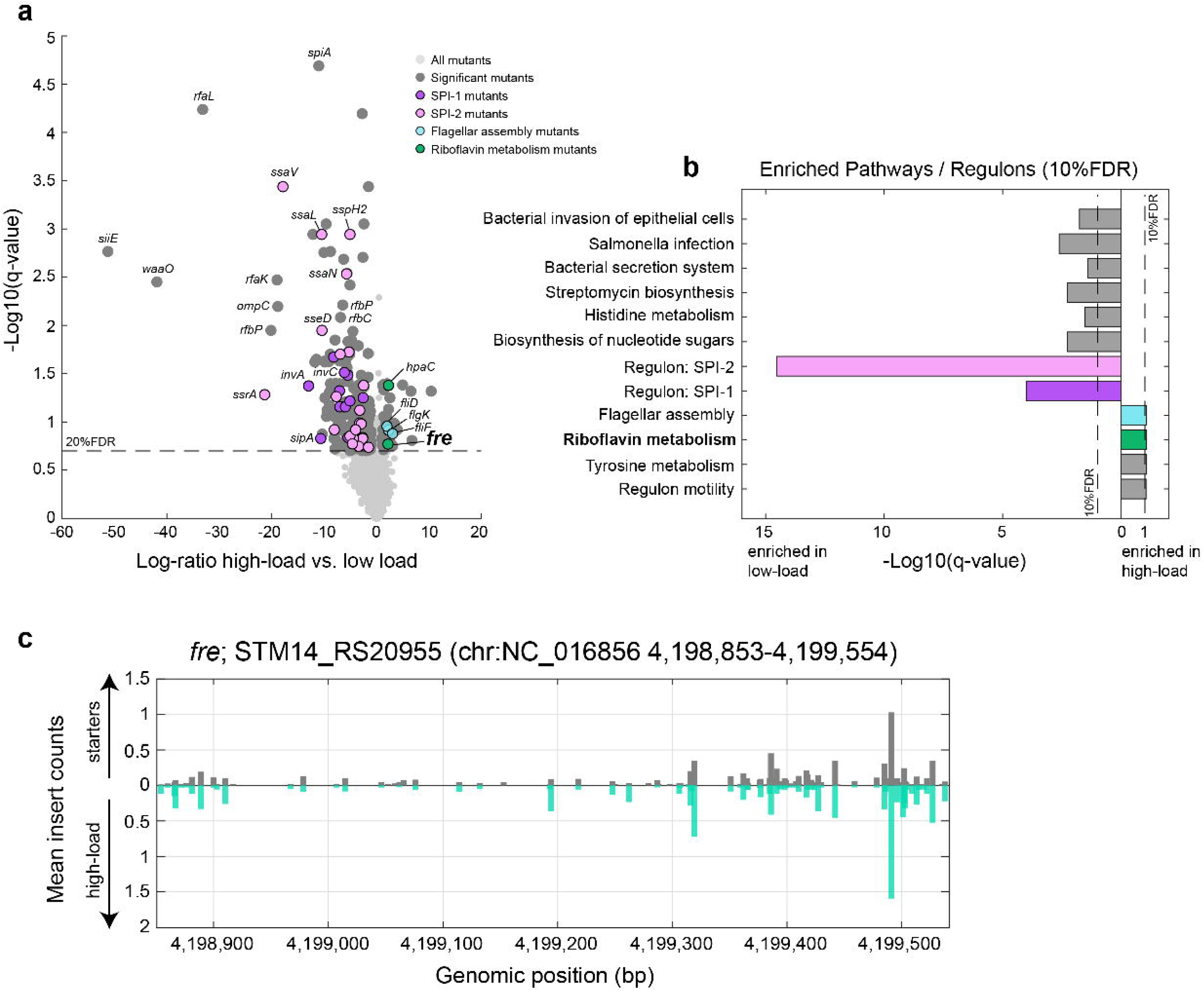
ICS-coupled transposon sequencing (nTraDIS) identifies bacterial determinants of intracellular replication. **a,** Transposon insertion sequencing of ICS-sorted macrophages identifies mutants enriched by intracellular bacterial load. Volcano plot of transposon-mutant enrichment between the high- and low-load gates (log-ratio high-load versus low-load) against significance (−log_10_ q-value); dashed line, 20% FDR. Mutants are colored by category (SPI-1, SPI-2, flagellar assembly, flavin/riboflavin metabolism), with selected genes labeled. **b,** The screen recovers known intracellular fitness pathways and identifies flavin metabolism. Pathway and regulon enrichment (10% FDR) among gate-enriched mutants; bars indicate enrichment in the low-load (left) or high-load (right) population. **c,** *fre*-disrupting insertions are selected for during intracellular replication. Transposon insertion profile across the fre locus (STM14_RS20955), showing mean insertion counts in the starter inoculum library (top) and the high-load gate (bottom) as a function of genomic position.

In the hyper-replicating population, mutants in flagellar assembly genes were enriched (*fliD*, *fliF*, *flgK*). Flagellin triggers NAIP/NLRC4-dependent caspase-1 activation and pyroptotic cell death of the infected macrophage [52]; loss of flagella therefore allows bacteria to evade this response and accumulate to high intracellular numbers, consistent with the attenuation of flagellin-overexpressing strains in vivo [53]. Remarkably, the high-load population was also enriched for transposon insertions in riboflavin metabolism, driven by two flavin reductase genes, *fre* and *hpaC*. The *fre* insertion profile showed marked enrichment across the gene body relative to the starter inoculum library (**Fig. 2c**), consistent with strong positive selection for *fre*-disrupting events during intracellular replication.

Together, these results establish ICS of infection phenotypes coupled with nTraDIS as a quantitative tool for dissecting bacterial fitness determinants at single-cell resolution directly from host cells. The convergence on two independent flavin reductase mutants (*fre* and *hpaC*) implicated flavin metabolism in restricting intracellular replication. Whereas HpaC is dedicated to flavin reduction within the 4-hydroxyphenylacetate catabolic pathway [54,55], Fre is the predominant cytosolic flavin reductase in Enterobacteriaceae and supplies reduced flavins to a wide range of downstream processes. In *E. coli*, Fre has previously been linked to oxidative stress [56–58]. We therefore focused our follow-up experiments on Fre as a candidate for a broad-acting determinant of intracellular fitness.

### *fre* mutant enhances *S.*Tm replication in macrophages

*fre* encodes a NAD(P)H:flavin oxidoreductase that supplies reduced flavins to a range of bacterial processes [59,60]. To validate the hyper-replicative phenotype detected in the screen, we generated a full knockout of *fre* and characterized its intracellular behavior in BMDM infections. We first confirmed that in LB broth Δ*fre* was not growing faster than WT, but in fact had a slightly longer doubling time, indicating that its increased intracellular replication does not stem from a general growth advantage (doubling times of 24.4 vs 21.5 min; **Fig. S3a,b**).

We next used three orthogonal assays to validate the hyper-replicative phenotype of Δ*fre* in macrophages. First, using imaging flow cytometry we measured similar numbers of invading bacteria at 1 hpi between WT and Δ*fre*. At 24 hpi, Δ*fre*-infected macrophages displayed a larger fraction of cells with higher bacterial loads (**Fig. 3a** and **S3c**). Second, using a TIMER reporter, which distinguishes replicating from non-growing bacteria based on differential maturation of its dTomato moiety [25], we measured a larger replicative subpopulation in Δ*fre*-infected macrophages compared to WT at 24 hpi (**Fig. 3b,c** and **S3d-f**). Third, Δ*fre* infections yielded higher intracellular colony-forming unit (CFU) over time than WT (Fig. 3d, left panel). Inducible expression of *fre* from a complementation plasmid restored WT-like replication (**Fig. 3d**, right panel), confirming that the phenotype is attributable to loss of *fre*.

**Fig. 3:**
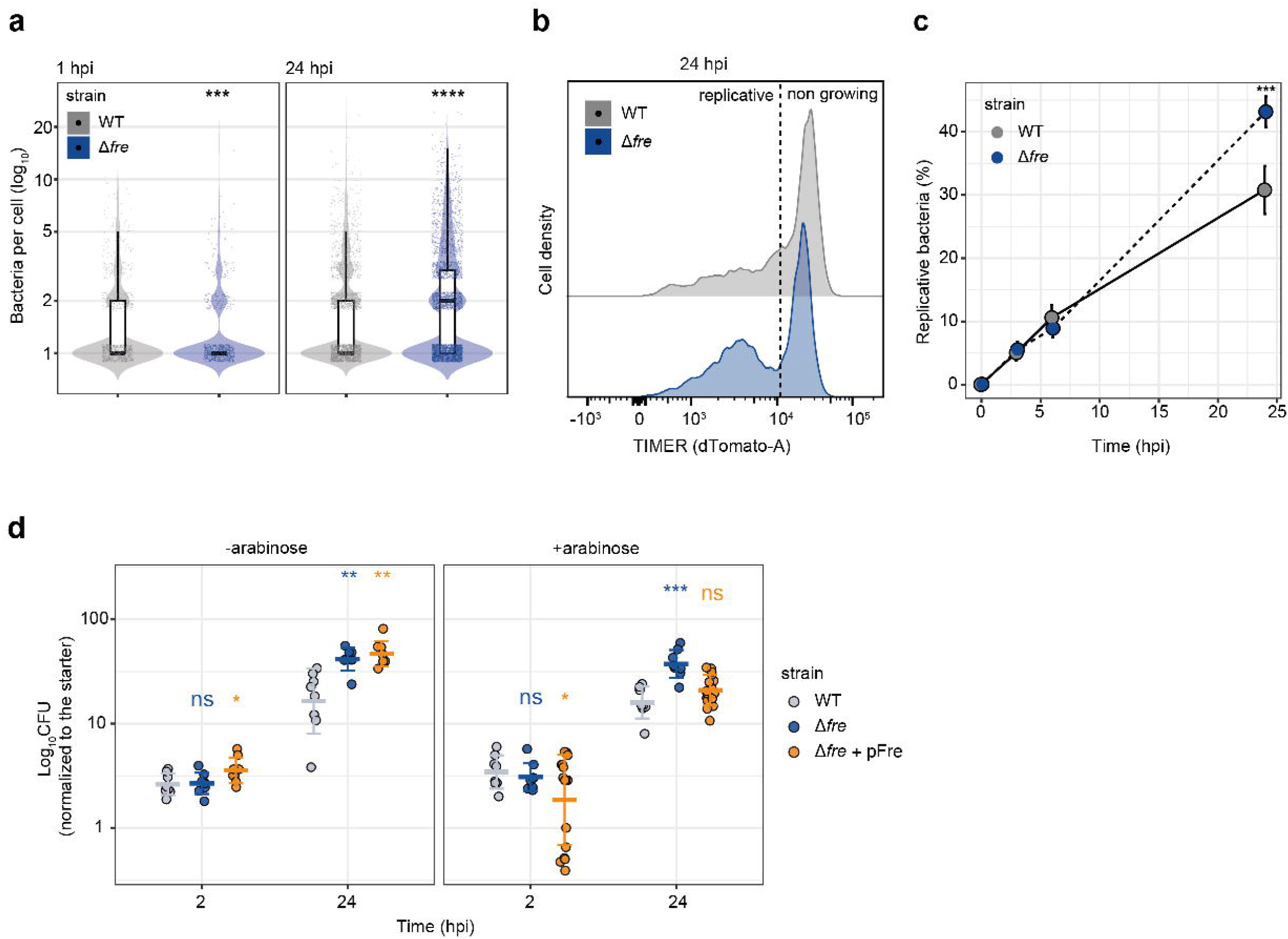
Loss of *fre* enhances *S.*Tm replication in macrophages. **a,** Δ*fre*-infected macrophages carry higher bacterial loads than WT at 24 hpi. Bacteria per cell measured by imaging flow cytometry (ImageStream) at 1 and 24 hpi for WT (grey) and Δ*fre* (blue); each point is one infected macrophage, bars indicate the median (n = 886 - 3261 cells per condition). **b,** Δ*fre* shows a larger replicative subpopulation than WT. TIMER (dTomato-A) fluorescence distributions at 24 hpi for WT (top) and Δfre (bottom); the dashed line separates replicative from non-growing bacteria. **c,** The replicative fraction increases over time and is greater in Δ*fre*. Percentage of replicative bacteria over the course of infection (0–24 hpi) for WT and Δ*fre*; points show mean ± SEM (t-test: ***, P ≤ 0.001 at 24 hpi). **d,** Complementation restores WT-like replication. Intracellular CFU normalized to the starter inoculum at 2 and 24 hpi for WT, Δ*fre*, and complemented Δ*fre* + pFre, without (left) and with (right) arabinose induction (n =8 – 16 independent infections). Significance markers are colored by strain (t-test: ns, P > 0.05; *, P ≤ 0.05; **, P ≤ 0.01; ***, P ≤ 0.001).

Finally, to test whether this phenotype extends to the in vivo setting, we performed a competition assay in a systemic mouse model by co-inoculating WT and Δ*fre* and measuring the competitive index in the spleen (**Fig. S3g**). Δ*fre* was outcompeted by WT in vivo, indicating that intracellular macrophage phenotype cannot fully account for the in vivo infection settings.

Together, these data show that loss of *fre* produces hyper-replication within isolated macrophages. We next sought to understand the mechanism by which *fre* restricts *S.*Tm replication in macrophages.

### Fre amplifies bacterial vulnerability to reactive oxygen species in vitro

Macrophages restrict intracellular *S.*Tm in part through reactive oxygen species (ROS), with hydrogen peroxide (H_2_O_2_) as one of the central effectors [61]. *S.*Tm in turn detoxifies H_2_O_2_ through a battery of partly redundant catalases and alkyl hydroperoxide reductases that contribute to its resistance to the oxidative burst and to virulence [7,62]. H_2_O_2_ damages bacterial macromolecules through multiple routes, one of which is the Fenton reaction, the iron-catalyzed conversion of H_2_O_2_ to the hydroxyl radical (•OH), which consumes Fe^2+^ and therefore depends on continuous regeneration of the ferrous iron pool [63,64]. In *E. coli*, reduced flavins supplied by Fre carry out this reduction of Fe^3+^ to Fe^2+^, and they become the dominant electron source for it when respiration is limited [57]. We therefore asked whether the same chemistry operates in *S.*Tm within the macrophage. If so, Fre should contribute to H_2_O_2_ toxicity in intracellular *S.*Tm, and its loss should attenuate oxidative damage.

To focus specifically on the H_2_O_2_ stress, we exposed WT and Δ*fre S.*Tm to a range of H_2_O_2_ concentrations in liquid culture. WT growth was strongly inhibited at 1 mM H_2_O_2_ and higher, whereas Δ*fre* sustained growth up to 5 mM with a concentration-dependent delay, indicating that loss of *fre* confers tolerance to H_2_O_2_ (**Fig. 4a**). Both strains were equally killed at 20 mM H_2_O_2_.

**Fig. 4:**
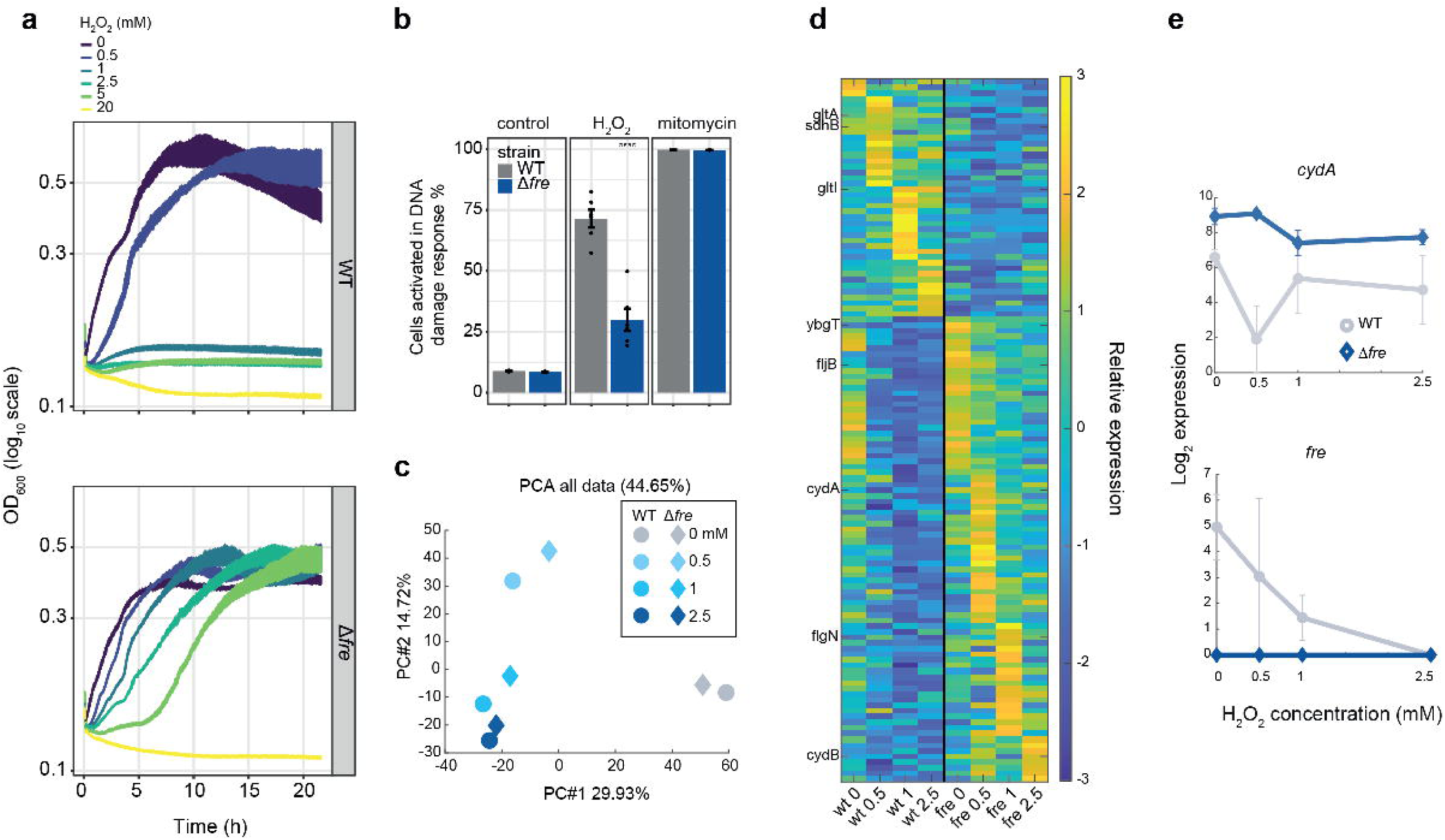
Fre amplifies *S.*Tm vulnerability to oxidative stress in vitro. **a,** Δ*fre* tolerates higher concentrations of H_2_O_2_ than WT. Growth (log_10_ OD_600_) of WT (upper panel) and Δ*fre* (lower panel) across a range of H_2_O_2_ concentrations (0–20 mM); shaded bands, mean ± SEM of 6 biological replicates. **b,** Δ*fre* sustains less H_2_O_2_-induced DNA damage than WT. Fraction of cells activating the SOS/DNA-damage reporter (pCdap-GFP) after 2 h exposure to 0.5 mM H_2_O_2_, 0.5 µg/ml mitomycin C, or no treatment, for WT (grey) and Δ*fre* (blue); mean ± SEM (n = 6 biological replicates; t-test: ****, P ≤ 0.0001). The DNA-crosslinking agent mitomycin C serves as a strain-independent positive control. **c,** H_2_O_2_ exposure is the dominant source of transcriptional variation. PCA of RNA-seq samples (PC1, 29.9%; PC2, 14.7%) for WT and Δ*fre* across H_2_O_2_ doses (averaged replicates shown). **d,** Loss of *fre* represses TCA-cycle genes while inducing motility and c*ydAB*. Heatmap of genes differentially expressed between WT and Δ*fre* across the H_2_O_2_ dose series (0, 0.5, 1, 2.5 mM), row-scaled relative expression; representative genes labeled. **e,** *fre* is transcriptionally repressed by H_2_O_2_ in WT, while the *cydAB* operon is constitutively elevated in Δ*fre*. Transcript levels (log_2_ expression) of *cydA* and *fre* as a function of H_2_O_2_ concentration in WT (grey) and Δ*fre* (blue); mean ± SD.

We next tested whether this tolerance reflected reduced oxidative damage to DNA. We introduced a fluorescent SOS reporter (p*Cda*p-GFP [65,66]), and measured activation in ∼70% of WT cells under H_2_O_2_ exposure, compared to ∼30% of Δ*fre* cells (**Fig. 4b**). As a control, the DNA-crosslinking agent mitomycin C activated the reporter in essentially all cells of both strains (**Fig. 4b**), indicating that the SOS response is intact in Δ*fre* and that the defect is specific to H_2_O_2_-induced damage.

To define the differences in adaptation of WT and Δ*fre* bacteria to peroxide stress, we performed RNA-seq across four H_2_O_2_ doses (0, 0.5, 1, and 2.5 mM). Principal component analysis (PCA) separated samples primarily by H_2_O_2_ presence along PC1 (29.9% variance) and by dose along PC2 (14.7% variance; **Fig. 4c**). Both genotypes shifted along PC1 upon H_2_O_2_ exposure, but at 0.5 and 1 mM Δ*fre* samples were displaced toward the untreated condition relative to dose-matched WT. At 2.5 mM the two genotypes converged. Hundreds of genes were induced or repressed by all H_2_O_2_ concentrations to comparable levels in both genotypes. OxyR targets emerged as the strongest induced regulon, while Fur targets were prominent among both induced and repressed gene sets (**Fig. 4d**). Canonical OxyR targets (*ahpC*, *ahpF*, *katG*, *dps*, *recA*) were induced to plateau levels across the dose series (**Fig. S4b**), indicating that direct H_2_O_2_ sensing through OxyR is intact in Δ*fre*.

The genotype effect therefore lies elsewhere in the response. Several genes were differentially expressed between WT and Δ*fre* at matched H_2_O_2_ doses (**Fig. 4d**). Genes repressed in Δ*fre* were most strongly enriched for the TCA cycle (*sdhAB*, *sucCD*, *gltA*, *acnA*; **Fig. S4c,d**), whereas genes induced in Δ*fre* were enriched for motility (*cheW*, *tsr*, *fljB*, *flgN*), ribosome and translation, and terminal respiration (*cydAB)* (**Fig. 4d** and **S4e**). *cydAB* was elevated in Δ*fre* across the entire dose series, including in untreated cultures (**Fig. 4e** and **Fig. S4f**). Loss of *fre* is thus accompanied by a reciprocal change in respiratory gene expression: reduced expression of the Fe–S-rich enzymes of the TCA cycle alongside increased expression of an Fe–S-independent terminal oxidase. Interestingly, in the WT response itself, *fre* was strongly and dose-dependently repressed by H_2_O_2_ (**Fig. 4e**, right panel), identifying *fre* transcription as a component of the WT peroxide response.

Together, these results show that Fre is not protective against peroxide but potentiates its toxicity: Δ*fre* sustained less DNA damage and growth inhibition than WT. This tolerance is accompanied by a transcriptional response in which WT represses *fre* upon peroxide exposure, while loss of *fre* leaves the OxyR antioxidant response intact but shifts respiratory gene expression toward cytochrome bd-I.

### **Δ***fre* protective phenotype in macrophages is mediated by cytochrome bd-I

Given Fre’s role in amplifying H_2_O_2_ toxicity in vitro, we asked whether the macrophage hyper-replication of Δ*fre* depends on host-derived ROS. We repeated the CFU assay in J774A.1 macrophages lacking the gp91^phox subunit of the NADPH oxidase (Δ*Cybb*), which cannot mount a respiratory burst [67]. Δ*fre* retained a ∼2-fold replication advantage over WT in Δ*Cybb* macrophages at 24 hpi, comparable to that observed in WT J774A.1 cells (**Fig. S5a,b**); inducible complementation rescued the phenotype in both. The macrophage advantage of Δ*fre* therefore does not require NADPH oxidase activity, indicating that the host respiratory burst is not the sole source of the oxidative stress to which Fre contributes. Consistent with this, an OxyR-dependent *katG* reporter in intracellular *S.*Tm was not induced above its baseline [68], indicating that intramacrophage bacteria do not sense substantial H_2_O_2_.

The persistence of the Δ*fre* advantage in Δ*Cybb* macrophages raised the question of which other antimicrobial effectors might contribute. Macrophages also produce nitric oxide (NO) via inducible nitric oxide synthase (iNOS) [69], and Fe–S cluster proteins, the same class showing dampened damage signatures in our RNA-seq, are well-documented targets of NO [70,71]. Moreover, cytochrome bd-I (*cydAB*), which we found upregulated in Δ*fre* during H_2_O_2_ treatment (**Fig. 4d,e**), is an NO-insensitive terminal oxidase that also scavenges ROS, allowing it to protect against both nitrosative and oxidative stress [72–76]. We therefore asked whether the tolerance of Δ*fre* extends to nitrosative stress. Using the NO donor NONOate, WT and Δ*fre* showed comparable dose-dependent growth delays at 1 and 5 mM (**Fig. 5a**). At 10 mM, both strains still grew but with a pronounced lag, which was substantially shorter in Δ*fre* than in WT (**Fig. 5b**). Loss of *fre* therefore confers accelerated recovery from NO stress, paralleling its protective effect against H_2_O_2_.

**Fig. 5:**
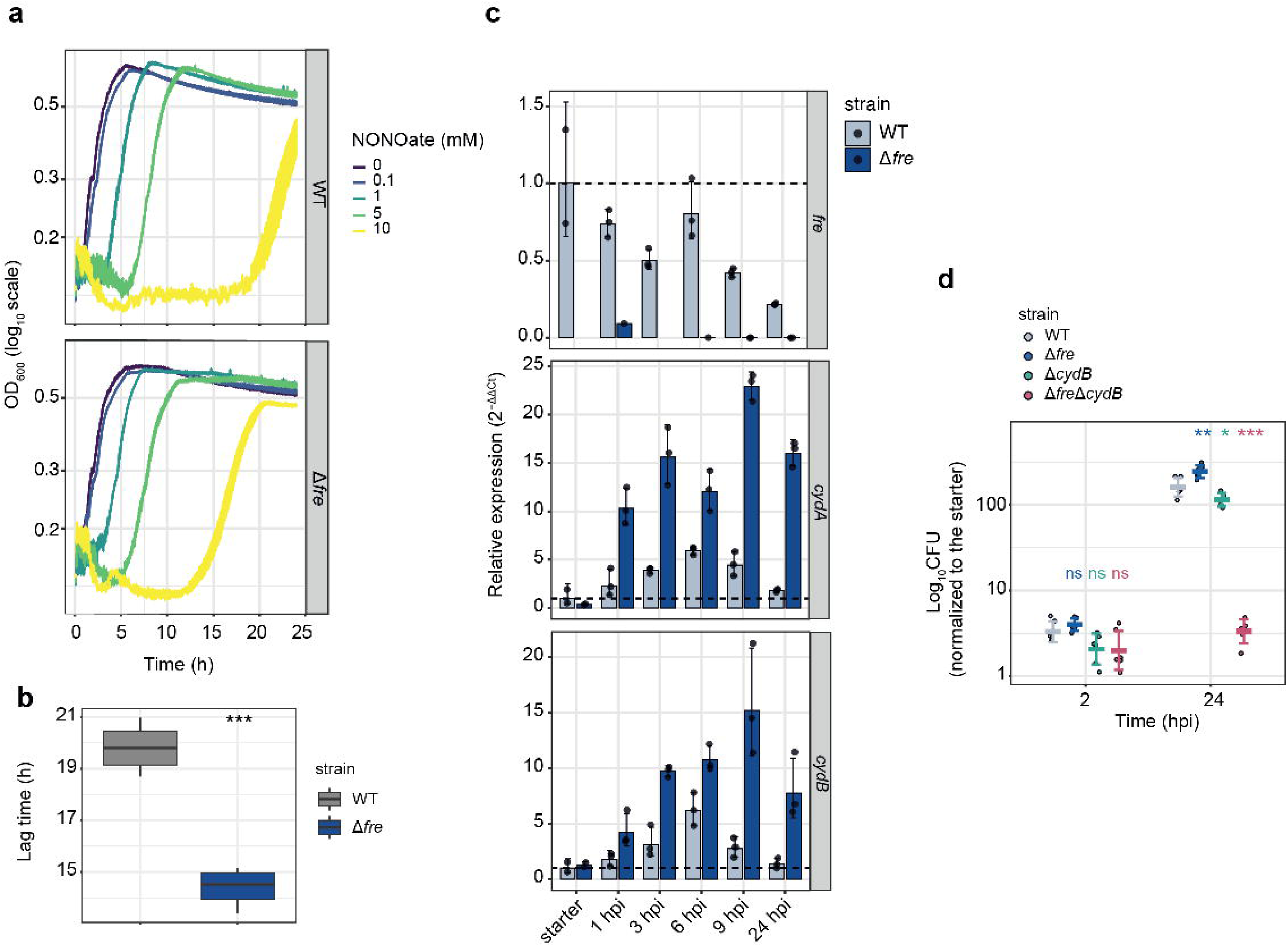
Δ*fre* protective phenotype in macrophages is mediated by cytochrome bd-I. **a,** Δ*fre* recovers faster from nitrosative stress than WT. Growth (log_10_ OD_600_) of WT (upper panel) and Δ*fre* (lower panel) across NONOate concentrations (0–10 mM), shaded bands, mean ± SEM of 4 biological replicates. **b,** Δ*fre* exits the NO-induced lag earlier than WT. Lag time extracted from the growth curves in Fig. 5a, at 10 mM NONOate for WT (grey) and Δ*fre* (blue) (n = 4; t-test: ***, P ≤ 0.001). **c,** RT-qPCR of *fre*, *cydA*, and *cydB* in wild-type (grey) and Δ*fre* (blue) *S.*Tm, from the starter inoculum and from bacteria recovered from infected J774A.1 at 1, 3, 6, 9, and 24 hpi. Transcript levels are expressed as 2^−ΔΔCt^ relative to the WT starter inoculum and normalized to *gmk*; the dashed line at 1 marks the WT starter baseline. *fre* confirms its infection-dependent repression in WT and its absence in Δ*fre*. *cydA* and *cydB* are progressively induced in Δ*fre* over the infection time course, exceeding WT at all intracellular time points; *cydA* and *cydB* are shown separately as independent amplicons for the same operon. Points, biological duplicates (for the starter) or triplicates (for the rest of the samples); bars, mean ± SD. **d,** Cytochrome bd is required for intracellular replication and for the Δ*fre* advantage. Intracellular CFU of WT, Δ*fre*, Δ*cydB*, and Δ*fre*Δ*cydB S.*Tm in J774A.1 at 2 and 24 hpi, normalized to the starter inoculum. All strains are comparable to WT at 2 hpi, whereas by 24 hpi Δ*fre* replicates above WT, Δ*cydB* slightly below it, and Δ*fre*Δ*cydB* drops to near-input levels, well below WT (n = 6). Points, individual infections; crossbar, mean; error bars, ±1 SD. Significance versus WT, colored by strain (ns, P > 0.05; *, P ≤ 0.05; **, P ≤ 0.01; ***, P ≤ 0.001).

To test whether this NO tolerance reflects *cydAB* activity and operates during infection, we first asked whether *cydAB* is induced within macrophages. RT-qPCR of macrophage-internalized bacteria confirmed that *cydAB* is upregulated in Δ*fre* relative to WT during infection (**Fig. 5c**), showing that the elevated *cydAB* expression observed in vitro is recapitulated within macrophages. To determine whether the Δ*fre* intracellular advantage requires this activity, we constructed a Δ*fre*Δ*cydB* double mutant. At 2 hpi, intracellular loads were comparable across all four strains (**Fig. 5d**, left panel), indicating equivalent uptake. By 24 hpi, Δ*fre* had replicated to higher levels than WT, confirming its intracellular growth advantage, whereas the Δ*cydB* single mutant showed only a modest reduction relative to WT (**Fig. 5d**, right panel). In contrast, the Δ*fre*Δ*cydB* double mutant failed to replicate intracellularly, remaining near the input level at 24 hpi, well below both WT and the Δ*cydB* single mutant. Thus, *cydB* is largely dispensable for intracellular replication in a WT background but becomes essential in the absence of *fre*, revealing a dependence in which Δ*fre* not only requires cytochrome bd-I for its proliferation advantage but relies on it for intracellular replication altogether.

Together, these results show that loss of *fre* protects *S.*Tm against a macrophage-derived stress distinct from the NADPH oxidase burst, and that this protection requires cytochrome bd-I. Given the insensitivity of bd-I to nitric oxide, and the tolerance of Δ*fre* to nitrosative stress in vitro, this stress may be nitrosative in origin.

## Discussion

Intracellular pathogens face distinct fates within their host: they may be killed, persist as non-replicating or slow-growing cells, or establish a replicative niche and expand. Single-cell approaches resolve these fates but not their genetic basis, whereas transposon screens assign genes to fitness while averaging across fates, leaving the bacterial functions that underlie each phenotype largely unresolved. Here, we developed an experimental approach that allows the study of genetic bacterial elements that impacts these phenotypes, by coupling transposon library screening to image-enabled cell sorting. Our approach provides accurate image-derived quantification of bacteria in a host cell at sort speeds compatible with functional screening. Applied to a *S.*Tm transposon library in BMDMs, sorting infected BMDMs by intracellular bacterial load allowed us to analyze bacterial mutants enriched within a defined population (here, the non-replicative and hyper-replicating cells). Beyond identifying genes required for replication, which conventional transposon screens already recover, this approach uncovered gene products that are not required for replication but instead restrain it, and whose loss increases bacterial load. Our results thus reveal that WT *S.*Tm carries functions that are counterproductive in the macrophage environment.

Two independent flavin reductases, *fre* and *hpaC*, were recovered in the hyper-replicating population, pointing to reduced flavins as a constraint on intracellular replication. We examined this through Fre, whose characterization indicated that its activity amplifies *S.*Tm vulnerability to both oxidative and nitrosative damage, with consequences that converge on iron–sulfur (Fe–S) cluster proteins. Three observations support this model. First, Δ*fre* bacteria tolerate sub-lethal doses of both H_2_O_2_ and the NO donor NONOate in vitro, two stresses whose toxicity is exerted in part through Fe–S cluster damage [70]. Second, the H_2_O_2_ transcriptional response in Δ*fre* shows reduced expression of TCA-cycle and respiratory enzymes (*sdhAB*, *sucCD*, *gltA*, *acnA*), several of which depend on Fe–S clusters, while the direct H_2_O_2_-sensing core (*ahpC*, *ahpF*, *katG*, *dps*, *recA*) is engaged normally. Third, Δ*fre* shows a concomitant upregulation of cytochrome bd-I, a terminal oxidase that lacks Fe–S clusters and is known to confer tolerance to both H_2_O_2_ and NO [75]. Deleting *cydB* eliminates the Δ*fre* replication advantage, showing that the induction of cytochrome bd-I is not only associated with the protective phenotype but necessary for it. A strikingly similar metabolic rewiring has previously been described in *E. coli*, where *fre* mutations suppress the lethality of hyper-initiation in Regulatory Inactivation of DnaA (RIDA)-deficient cells by rerouting electron flow toward cytochrome bd-I, which acts as an electron sink that lowers reactive oxygen species formation and consequent oxidative damage to DNA [76]. That a comparable rewiring is engaged in *S.*Tm Δ*fre* during both peroxide response and macrophage infection suggests it represents a general metabolic adaptation to loss of cytosolic flavin reduction. Consistent with the established role of cytochrome bd-I in NO protection [72,74], and with the persistence of the Δ*fre* advantage in Δ*Cybb* macrophages lacking a functional NADPH oxidase, these observations point to nitric oxide as a principal macrophage-derived stressor against which loss of *fre* is protective.

The opposite phenotype of Δ*fre* during systemic infection indicates that the same activity that restrains *S.*Tm replication in macrophages is necessary elsewhere in the infection cycle. This systemic disadvantage may be explained in part by the intrinsic growth cost of losing *fre*: the slower doubling we observed in LB compounds into a competitive disadvantage over successive generations that would contribute to the reduced spleen competitive index. An additional possibility is that *fre* is specifically required to counter the antimicrobial chemistries of other phagocytes. In vivo, *S.*Tm encounters multiple phagocyte types, including neutrophils, inflammatory monocytes, and dendritic cells, which differ in the antimicrobial chemistries they deploy [27]. Neutrophils, in particular, produce hypochlorous acid (HOCl) through the myeloperoxidase system, which oxidizes methionine residues in periplasmic proteins. Repair of this damage is mediated by the periplasmic methionine sulfoxide reductase MsrPQ [77], whose membrane component MsrQ has been shown to receive electrons from Fre [78,79]. Loss of *fre* could therefore compromise repair of HOCl-induced damage, even while remaining advantageous in the H_2_O_2_-rich environment of the macrophage phagosome. Together, these considerations suggest that the systemic cost of losing *fre* may combine an intrinsic growth disadvantage with a cell-type-specific requirement, such that the net consequence of Fre activity for *S.*Tm fitness depends in part on which antimicrobial strategy a given host cell employs.

In summary, we have shown that image-enabled cell sorting of infected macrophages allows the identification of bacterial determinants of intracellular replication, and have used this approach to uncover a previously unrecognized role for the flavin reductase Fre as a context-dependent restraint on *S.*Tm replication. By coupling flavin reduction to oxidative and nitrosative damage, Fre amplifies the toxicity of multiple antimicrobial chemistries. Its loss shifts respiratory gene expression toward the protective cytochrome bd-I oxidase, an adaptation that is advantageous within macrophages but detrimental elsewhere in the infection cycle. Such context-dependent trade-offs may be a common feature of the intracellular lifestyle, where pathogens transit between cellular environments imposing distinct chemistries of attack. Understanding how *S.*Tm and other intracellular pathogens navigate distinct selective pressures will help define the bacterial trade-offs that shape the outcome of infection.

Our approach can further provide a framework for extending this dissection from the pathogen to the host, for example, by coupling it to a host-directed CRISPR/Cas9 library, while single-cell or dual RNA-seq of sorted cells would report host and bacterial transcriptomes from the same phenotypic class. More broadly, any infection phenotype that can be captured in an image becomes a sortable variable, allowing both sides of a divergent outcome to be dissected within a single framework.

## Methods

### Bacterial strains

All strains used in this study are listed in Supplementary Table 1.

### Bacterial growth conditions

*S.* Typhimurium strain 14028S and derivatives were routinely grown in lysogeny broth (LB) at 37°C with shaking at 220 rpm, supplemented with antibiotics as appropriate (kanamycin 50 µg/ml; chloramphenicol 30 µg/ml; ampicillin 100 µg/ml).

### Lambda-red recombination

The *fre* and *cydB* genes were deleted in *S.* Typhimurium 14028S by λ-Red recombineering as described in [80]. Briefly, a kanamycin or chloramphenicol resistance cassette flanked by FRT sites was amplified from pKD4 or pKD3, respectively, using primers carrying 40-bp homology arms to the regions flanking each target gene (fre_KO_FW/fre_KO_REV for *fre*; cydB_KO_FW/cydB_KO_REV for *cydB*; Supplementary Table S1). Each PCR product was electroporated into *S.* Typhimurium 14028S harboring pKD46, which expresses the λ-Red recombinase under arabinose induction. Transformants were selected on LB agar containing kanamycin (50 µg/ml) or chloramphenicol (30 µg/ml) at 37°C. Deletions were confirmed by colony PCR: for *fre*, using fre_KO_validation_FW paired with Cm_KO_REV (chloramphenicol-resistant mutants) or Kan_KO_REV (kanamycin-resistant mutants); for *cydB*, using cydB_KO_validation paired with Kan_KO_REV (Supplementary Table S1).

### Tn7-mediated chromosomal labelling of S. Typhimurium with beRFP

To obtain stable, single-copy chromosomal expression of beRFP, S. Typhimurium 14028S was labelled at the attTn7 site by triparental mating as previously described [81], with minor modifications. The DAP-auxotrophic donor strain *E. coli* MFDpir was transformed with either the Tn7 transposase helper plasmid pJMP1039 (Addgene, #119239) or a Tn7 transposon delivery plasmid derived from pJMP1339 (Addgene, #119271) in which mBeRFP was placed under the expression of an artificial promoter [82] (Addgene #260343). Donor strains were grown overnight at 37°C in LB supplemented with 300 µM DAP (Sigma, D1377) and 100 µg/ml ampicillin. The *S.*Tm recipient was grown in parallel for ∼16 h in LB at 37°C without antibiotics. For each mating, 100 µl of each donor strain and 100 µl of recipient were combined in 700 µl LB, pelleted (2 min, 7,000 × g), washed twice in 1 ml LB, resuspended in 30 µl LB, spotted onto an LB plate, and incubated overnight at 30°C. Mating spots were resuspended in 100 µl PBS and plated onto LB agar containing kanamycin (50 µg/ml) and lacking DAP, selecting simultaneously for chromosomal insertion of the beRFP cassette in S. Typhimurium and against the MFDpir donor. Insertion at attTn7 was verified by colony PCR using primers listed in Supplementary Table S1. The kanamycin resistance cassette was subsequently resolved by transformation with the temperature-sensitive Flp recombinase plasmid pCP20 [83] by curing at 42 °C, with loss of both the cassette and pCP20 confirmed by replica plating and PCR. Verified strains were stored at −80°C in LB + 20% glycerol.

### Construction of fluorescent reporter and complementation vectors

The plasmid pCdap-GFP (Addgene #260344) was assembled by Gibson assembly using pSC101 as the backbone. pSC101 was linearized by digestion with XhoI (NEB, R0146S) and XbaI-HF (NEB, R0145S). Insert 1 encoding the *cda* promoter was a synthesized DNA fragment (Twist Bioscience), and insert 2 (GFP) was amplified from pFPV25 (Addgene #20667) plasmid using primers GFP_FW and GFP_REV (Supplementary Table S1). The six-color reporter constructs were generated by ordering codon-optimized sequences for mRuby (Addgene #260345), mVenus (Addgene #260346), cyOFP (Addgene #260347), mBeRFP (Addgene #260348), mKalama1 (Addgene #260349), and mKate2 (Addgene #260350) as gene fragments (Twist Bioscience), each preceded by a synthetic promoter [82], and cloning them by Gibson assembly into pFPV25 (Addgene #20667) from which the GFP cassette had been removed. For *fre* complementation, the full-length *fre* coding sequence was amplified from *S.*Tm 14028S genomic DNA using primers fre_compl_FW and fre_compl_REV (Supplementary Table S1) and cloned by Gibson assembly into pBAD24 downstream of the arabinose-inducible PBAD promoter. The resulting plasmid (pFre, Addgene #260351) was introduced into Δ*fre* by electroporation. For infection experiments, complemented strains were induced with arabinose directly in DMEM at the time of infection. All plasmid sequences were confirmed by Sanger sequencing.

### BMDM isolation

Bone marrow was extracted from mice as previously described[84], with some modifications. Female C57BL/6 mice, aged 8 weeks, were euthanized and femurs and tibias were harvested. Bone marrow was flushed from the bones, resuspended in DMEM, and plated in tissue culture treated dishes (Corning, 430599) in DMEM (Thermo Fisher, 41965-039) medium containing 1 mM sodium pyruvate (Biological Industries, 03-042-1B), 20% fetal bovine serum (FBS) (Thermo Fisher, 10270106), and 20 ng/ml recombinant mouse M-CSF (rmM-CSF, BioLegend, 576408). Cells were harvested at day 6 after bone marrow isolation and frozen until use. All bone marrow extractions were performed in accordance with the Institutional Animal Care and Use Committee at the Weizmann Institute (IACUC Approval number 04520725-1).

### BMDM infection

BMDMs (8x10^5^ cells/well) were plated in untreated six–well plates (Greiner, 657185) supplemented with DMEM (Thermo Fisher, 41965-039) medium containing 1 mM sodium pyruvate (Biological Industries, 03-042-1B), 20% fetal bovine serum (FBS) (Thermo Fisher, 10270106), and 20 ng/ml recombinant mouse M-CSF (rmM-CSF, BioLegend, 576408). Overnight LB-grown cultures of *S*.Tm were washed with 2x PBS before infection. BMDMs were infected at a multiplicity of infection (MOI) of 2:1 unless otherwise specified, and spun down for 5 min at 400*xg* to synchronize internalization. After 30 minutes, cells were washed with medium containing 50 *µ*g/ml gentamicin to remove non-internalized *S*.Tm. Fresh medium containing 50 *µ*g/ml gentamicin was then added back to the cells for the duration of infection.

### Cell harvesting, fixation and immunostaining

At each time point, infected BMDMs were washed once with DPBS -/- (Biological Industries, 02-023-1A) and detached by incubation in 300 µl of DPBS -/- supplemented with 5 mM EDTA for 5 min on ice. Cells were pelleted (5 min, 400 × g, 4°C) and fixed/permeabilized in ice-cold 80% methanol for 15 min on ice. After two washes in FACS buffer (PBS + 1% BSA, 5 mM EDTA), cells were stained for 1 h at 4°C with a FITC-conjugated anti-Salmonella O:4 antibody (clone 1E6, Santa Cruz sc-52223) at 50 ng per 10^6^ cells, washed three times in FACS buffer, and analyzed/sorted by ICS or ImageStream.

### Image-enabled cell sorting (ICS)

ICS was performed on a prototype of the BD FACSDiscover™ S8 sorter equipped with BD CellView™ Image Technology (BD Biosciences) as described [36]. Sorts used a 100 µm nozzle and a sheath pressure of 20 psi, in purity mode, with the input and collection sample chambers held at 4°C. Cells were first gated on FSC-A × SSC-A, followed by sequential doublet exclusion on SSC-W × SSC-H and FSC-W × FSC-H, and a final image-based singlet gate using radial moment versus eccentricity of the light-loss signal. Gates were iteratively refined during acquisition by visual inspection of single-cell images. The gating strategy using a combination of traditional signal intensity parameters and image-derived parameters is shown in Fig. 1a and S1a. In brief, infected (LPS+) cells were then partitioned by bacterial load on FITC-channel image features. The low-load gate (1 bacterium per host cell) was defined on FITC-A versus FITC diffusivity and the high-load gate (≥5 bacteria per host cell) on FITC size versus FITC max intensity. FITC size is the number of pixels above a user-defined threshold, FITC max intensity is the value of the brightest pixel, FITC-A is the integrated fluorescence signal (area) of the event, and FITC diffusivity measures how spread the signal is across the event, with low values for a concentrated punctum and high values for a signal distributed over a larger area. Sorted populations were collected into PBS supplemented with 3% BSA.

### Confocal Microscopy and validation of ICS sorting

BMDMs were sorted with ICS and imaged on a Zeiss LSM780 confocal microscope using a 40×/1.4 NA Oil DIC Plan-Apochromat objective (Zeiss). Image analysis was performed in Fiji. Intracellular bacteria were identified from the LPS (FITC) channel by thresholding and Analyze Particles, yielding the x,y coordinates of each bacterium. Host-cell boundaries were segmented from the brightfield channel using rajlabimagetools segmentation pipeline (https://github.com/arjunrajlaboratory/rajlabimagetools/wiki). Bacteria were then assigned to host cells in MATLAB: each cell mask was converted to a polygon and the number of bacterial coordinates falling within each polygon was counted (point-in-polygon test), giving the number of bacteria per host cell.

### Flow cytometry

At the indicated time points, infected BMDMs were washed once with DPBS -/- and lysed in 0.1% Triton X-100 in PBS for 5 minutes at room temperature to release intracellular bacteria. Lysates were diluted into FACS buffer before acquisition. Bacterial samples were acquired on a BD FACSAria III (BD Biosciences) fitted with a 70 µm nozzle. Data were analysed with FlowJo v10 (BD Biosciences).

### ImageStream analysis

Quantification of intracellular bacterial load in WT and Δ*fre* infections was performed on an Amnis ImageStream®X Mark II (Cytek). BMDMs were infected, fixed, and immunostained with anti- *Salmonella* O:4 FITC as described above. Single cells were gated using the Area_M01 × Aspect Ratio_M01 features, followed by focus selection, and infected cells were identified as FITC-positive. Bacterial load per cell was quantified using the Spot Count feature on the FITC channel. Analysis was performed in IDEAS software.

### CFUs

For intracellular CFU, infected BMDMs were washed once with DPBS -/- and lysed in 0.1% Triton X-100 in PBS for 5 minutes at room temperature. For in vivo CFU, spleens were homogenized in 5 ml of homogenization buffer (PBS, 5% FBS, 10 mM EDTA). Lysates/homogenates were 10-fold serially diluted in PBS and plated onto LB agar plates containing the appropriate antibiotic. Plates were incubated overnight at 37°C and colonies counted using a Scan® 500 automatic colony counter (Interscience). For competitive index experiments, WT and Δfre were distinguished by plating on LB ± kanamycin. The competitive index (CI) was calculated as the ratio of mutant to WT CFU.

### Growth curves and doubling-time analysis

Overnight LB cultures of wild-type and Δ*fre S.*Tm were diluted 1:1000 into fresh LB, dispensed into a 96-well plate, and grown at 37°C with agitation in a plate reader (Cytation 5, Agilent). OD_600_ was recorded every 5 min for 16 h. Growth parameters were extracted in R using `fit_easylinear()` from the `growthrates` package, which fits a linear model to log-transformed OD over the steepest-slope window of consecutive timepoints (h = 20, quota = 0.95). Prior to fitting, each well was baseline-corrected by subtracting its minimum OD_600_. The maximum specific growth rate (µmax) was taken as the slope of the fit, and doubling time was calculated as ln(2)/µmax. Data are from 24 biological replicates per strain. Doubling times were compared between strains by two-sided Student’s t-test.

### SOS response assay

DNA damage was quantified using the SOS reporter plasmid pCdap-GFP (see vector construction), in which gfp is expressed from the LexA-regulated cda promoter. WT and Δ*fre S.*Tm carrying pCdap-GFP were grown in M9 minimal medium as above to OD_600_ 0.4, then exposed to 0.5 mM H_2_O_2_ (Sigma, 88597-100ML-F) for 2 h at 37°C with shaking. As a positive control, parallel cultures were treated with 0.5 µg/ml mitomycin C (Roche, #10107409001) for 2 h. Untreated cultures served as the baseline. After treatment, cells were kept on ice and GFP fluorescence was measured by flow cytometry on a BD FACSAria III (BD Biosciences) with a 70 µm nozzle. The fraction of reporter-positive cells was determined by gating on GFP fluorescence relative to an untreated control. Data were analyzed in FlowJo v10 (BD Biosciences).

### M9 growth curves in H_2_O_2_ and NONOate

Wild-type and Δ*fre S.*Tm were grown in M9 minimal medium supplemented with 0.2% glycerol, 1 mM MgSO_4_, 200 µM CaCl_2_, 1 µg/ml thiamine, and 1 mg/ml casamino acids at 37°C for ∼3 h with shaking. The culture was then diluted to an OD_600_ of 0.375, and 100 μl aliquots were dispensed into a 96-well plate and treated with 0, 0.5, 1, 2.5, 5, or 20 mM H_2_O_2_ (Sigma, 88597-100ML-F) or with 0, 0.1, 1, 5, or 10 mM NONOate (Cayman Chemical Company, #82150). Plates were incubated for 24 h at 37°C with agitation in a multimode imaging reader (Cytation 5, Agilent), and growth was monitored by measuring absorbance at 600 nm every 5 min. Growth parameters were extracted in R using `fit_easylinear()` from the `growthrates` package as described for the LB growth curves. Lag time was taken as the lag coefficient returned by the fit.

### *In vivo* infection

All animal experiments were approved by the Institutional Animal Care and Use Committee of the Weizmann Institute of Science (IACUC approval number 00100124-2). Female C57BL/6 mice, 8–10 weeks old, were used. For competitive infections, wild-type and Δ*fre S.*Tm were grown overnight in LB, washed twice in PBS, and mixed at a 1:1 ratio. Mice were inoculated intravenously via the tail vein with 1,000 total CFU in 200 µl PBS. The exact input ratio was determined by plating serial dilutions on LB and LB + kanamycin. At 72 h post-infection, mice were euthanized and spleens were aseptically removed, homogenized in 5 ml of homogenization buffer (PBS, 5% FBS, 10 mM EDTA), passed through a 70-µm filter, serially diluted, and plated on LB and LB + kanamycin to enumerate total and Δ*fre* CFU, respectively. Wild-type CFU were calculated by subtracting Δ*fre* (kanamycin-resistant) counts from total counts. The competitive index (CI) is defined as the ratio of mutant to WT.

### Transposon library construction

Transposon library of *Salmonella* was generated using EZ-Tn5 <KAN-2>Tnp Transposome kit (Lucigen, TSM99K2) following manufacturer’s instructions. Briefly, 300 ml of *S*.Tm was grown in LB at 37°C to a final OD_600_ of 0.8. Cells were washed twice in 180 ml of ice-cold wash solution (300 mM sucrose), and then resuspended in 1 ml of wash solution. Each 100 *µ*l aliquot was transformed with 0.1 pmol of EZ-Tn5 <KAN-2>Tnp Transposome using 2 mm electroporation cuvettes (Cell Projects, EP-102) and electroporation program Ec2 (V = 2.5 kV) with MicroPulser Electroporator (Bio-rad). Each reaction was allowed to recover in 500 *µ*l SOC (2% tryptone, 0.5% yeast extract, 10 mM NaCl, 2.5 mM KCl, 10 mM MgCl_2_, 10 mM MgSO_4_, 20 mM glucose) for 45 minutes at 37°C, and then plated in a 15 cm LB agar plate supplemented with kanamycin 30 *µ*g/ml. On the following day transformants were scraped off of the LB plates and pooled into a library of approximately 250,000 transformants.

### nTraDIS sample preparation

DNA was extracted from a range of 350,000 - 1,300,000 sorted infected cells using DNeasy Blood & Tissue (Qiagen, 69504) following manufacturer’s instructions, with an overnight proteinase K digestion step, and resuspension in 200 *µ*l of resuspension buffer. DNA (∼ 400 - 1000 ng per sample) was processed as previously described [85], with few modifications. Briefly, DNA was sonicated at 4°C (program: 6 cycles of 30 s on, 60 s off) on a Bioruptor Plus (Diagenode), achieving a size distribution between 200-1000 nucleotides (nt), with 450 nt average size. The sonicated DNA was end-repaired for 30 minutes at 20°C in 200 *µ*l final volume containing 1x end-repair buffer and 10 *µ*l of end-repair mix (NEB, E6050L). The reaction was cleaned in 2x AmpureXP Beads (Beckman Coulter) and eluted in 37.5 *µ*l. A-tailing was performed 20 min at 37°C in 50 *µ*l final volume, containing 1x NEBuffer 2 (NEB, B7002S), 1 mM ATP, and 125U of Klenow fragment (NEB, M0212M). The reaction was cleaned in 2x AmpureXP Beads (Beckman Coulter) and eluted in 44 *µ*l. DNA was then dephosphorylated overnight at 37°C in 50 *µ*l final volume, containing 1x rCutSmart buffer (NEB, B6004S), and 5U of quick CIP (NEB, M0525L). The day after, the reaction was heat-inactivated for 2 minutes at 80°C, cleaned in 2x AmpureXP Beads (Beckman Coulter), and eluted in 18 *µ*l. The DNA template was ligated for 20 min at 25°C in 50 *µ*l final volume, containing 0.22 *µ*M Illumina-compatible forked indexed adapters, 1x buffer, and 5 *µ*l of quick ligase (NEB M2200L). To release the non-ligated strand of the adaptor, DNA was denatured at 96°C for 5 min, cleaned with 0.8x AmpureXP beads (Beckman Coulter), and eluted in 30 *µ*l. Two PCR reactions were used for TIS library construction. The first PCR reaction was performed in 50 *µ*l final volume containing 0.5 *µ*M TN bait primer 1 (Supplementary Table S1), 0.5 *µ*M Illumina enrichment primer, 1x KAPA HiFi HotStart ReadyMix (Roche, KK2601) and 10 *µ*l DNA template, using ∼ 150 - 200 ng as template and a number of reactions according to the divided input. PCR program: 60s 98°C, 15 cycles of 15s 98°C, 20s 68°C, 60s 72°C, and final extension of 60s 72°C. The first PCRs were pooled and cleaned with 1x AmpureXP beads (Beckman Coulter), and the eluate was used as a template in the second PCR reaction. Conditions for the second PCR reaction were identical to the first, with the following differences: reaction was performed in two 50 *µ*l reactions per sample and TN bait primer mix 2 consisted in a mix of staggered nested primers (Supplementary Table S1). Libraries were quantified with NEBNext Library Quant Kit for Illumina (NEB, E7630L) and sequenced on a NovaSeq 6000 machine with the following settings: R1=85, I1=8, and R2=37.

### nTraDIS analysis

Read1 was used to extract reads mapping to the transposon region, by searching for the sequence TTGAGATGTGTATAAGAGACAG (2 mismatches allowed). The region matching to the transposon was trimmed from R1 before mapping, only reads longer than 20 nt were kept. Transposon mapping R1 and R2 reads were then mapped to the *S*.Tm 14028S genome with the program bowtie, multimappers were excluded from the analysis. Read 1 was used to identify the disrupted gene. The random sonication point contained in read 2, was binned using a window size of 5 to remove mapping errors and overestimation of sonication points. The binned sonication points were used as a unique molecular identifier (UMI) to collapse PCR duplicates. Molecules with less than 3 read coverage were removed from the analysis. After UMI-based deduplication and read filtering, an average of 57.46 integrations per kilobase was recovered in coding sequences (CDSs) (total insert in CDS divided by the total length of the CDS). All samples were normalized to the number of inserts per sample by dividing the total insert count of each sample by the median total insert count across all samples in the experiment. The normalized data were then log2-transformed. The three starter inoculum samples were averaged, and log-ratio values were calculated for each insert in each sample relative to the starter average. To calculate a gene-level enrichment score relative to the starters, all positive log-ratios for inserts within each gene were summed. Finally, scores from the low-load and high-load samples were compared to identify mutants enriched and depleted in each population using a linear mixed-effects (LME) model, with condition (high/low-load) as a fixed effect and infection replicate as a random effect.

### RNA-seq sample preparation

Wild-type and Δ*fre S.*Tm were grown in M9 minimal medium supplemented with 0.2% glycerol, 1 mM MgSO_4_, 200 µM CaCl_2_, 1 µg ml^⁻^¹ thiamine, and 1 mg ml^⁻^¹ casamino acids at 37°C for ∼3 h with shaking. The culture was then diluted to an OD_600_ of 0.375, and 40-ml aliquots were treated with 0, 0.5, 1, or 2.5 mM H_2_O_2_ (Sigma, 88597-100ML-F) for 30 min at 37°C with shaking. Bacteria were harvested by centrifugation (7,000 × g, 5 min, RT) and snap-frozen in liquid nitrogen. Pellets were resuspended in 700 µl TRIzol (Thermo Fisher) and lysed by gentle bead-beating using an OmniLyse device (Claremont BioSolutions). Total RNA was extracted with the miRNeasy Mini Kit (QIAGEN), including on-column DNase I digestion followed by an additional treatment with TURBO DNase (Thermo Fisher). Strand-specific cDNA libraries were prepared using the RNAtag-seq protocol as previously described [86] and sequenced on an Illumina NovaSeq instrument.

### RNA-seq analysis of WT and **Δ***fre* bacteria in response to H_2_O_2_

FASTQ reads were demultiplexed using fastq-multx and aligned to the *S*.Tm 14028S genome using STAR (version 2.7.9). Gene counts were quantified from R2 reads using HTSeq. Count data were normalized using a library size factor calculated by dividing the total number of reads in each sample by the median of the total read count across all samples. Normalized counts were log2-transformed, and a minimum expression threshold of 3 was applied. Biological replicates were averaged, except for one WT replicate treated with 0.5 mM H_2_O_2_ (replicate #2), and one Δ*fre* replicate treated with 0.5 mM H_2_O_2_ (replicate #2), which were excluded from the analysis because they were deviated from the other three replicates. To identify genes differentially expressed in response to H_2_O_2_ in WT versus Δ*fre* bacteria, we calculated the differences in the integrals of each gene in Δ*fre* relative to WT at elevated H_2_O_2_ concentrations. The distribution of differences across all genes was approximately normal, with a mean of 0.3 and a standard deviation of 1.9. Genes with differences greater than 2 standard deviations from the mean were defined as differentially expressed between WT and Δ*fre* in response to H_2_O_2_.

### Generation of **Δ***Cybb* J774A.1 macrophages

The NADPH oxidase–deficient Δ*Cybb* line was generated from J774A.1 macrophages by CRISPR–Cas9 ribonucleoprotein (RNP) electroporation. Two Cybb-targeting crRNAs directed against the first exon were designed using the IDT CRISPR design tool (target sequences in Supplementary Table S1) and annealed with tracrRNA (IDT) to form crRNA:tracrRNA duplexes. RNP complexes were assembled by incubating the duplexes with Alt-R S.p. Cas9 Nuclease V3 (IDT) at room temperature for 10–20 min. For each reaction, 1 × 10^6^ cells were washed twice in Ca^2+^/Mg^2+^-free PBS, resuspended in supplemented P3 Nucleofector solution (Lonza SF Cell Line 4D-Nucleofector X Kit L), combined with the RNP complex and Alt-R Cas9 Electroporation Enhancer (IDT), and electroporated on a 4D-Nucleofector System (Lonza) using program CM-139. Cells were recovered in pre-warmed medium and allowed to expand for 48–72 h before clonal isolation by limiting dilution (∼ 0.1 cells/well) in 96-well plates. Single-cell clones were screened by PCR across the target site, and biallelic disruption of Cybb was confirmed by Sanger sequencing and by qPCR. A validated homozygous knockout clone was used for all subsequent experiments.

### RNA extraction and RT-qPCR

#### Bacterial gene expression

J774A.1 macrophages infected with *S.*Tm 14028S strains (WT, Δ*<u>fre</u>*) at MOI of 2:1 were harvested at 1, 3, 6, 9, and 24 hpi in TRIzol (Invitrogen/Thermo Fisher Scientific, Waltham, MA, USA). Total RNA was extracted using Direct-zol RNA MiniPrep (Zymo Research, Irvine, CA, USA) with on-column DNase I treatment, followed by a TURBO DNase treatment (Invitrogen/Thermo Fisher Scientific). cDNA was synthesized from 1.5 µg of total RNA using SuperScript IV (Invitrogen/Thermo Fisher Scientific) with random hexamer priming; no-reverse-transcriptase (–RT) controls were included to confirm the absence of genomic DNA. Quantitative PCR was performed on a CFX Opus 384 (Bio-Rad, Hercules, CA, USA) using iTaq Universal SYBR Green Supermix (Bio-Rad) with gene-specific primers (Supplementary Table S1). Cycling conditions were 95°C for 30 s, then 40 cycles of 95°C for 3 s and 60°C for 20 s, followed by melt-curve analysis to verify amplicon specificity. Transcript levels were normalized to the reference gene *gmk* and expressed relative to the WT reference condition using the 2^−ΔΔCt^ method[87]. Reactions were run in technical duplicates across two (for the starters) or three (for the rest) independent biological replicates.

#### Host gene expression

For host NADPH oxidase expression, total RNA from WT and Δ*Cybb* J774A.1 macrophages was extracted using the RNeasy Mini Kit (QIAGEN, Hilden, Germany) with on-column DNase I treatment. cDNA synthesis and qPCR were carried out as described for bacterial samples, using primers specific for murine *Cybb* and *Gapdh* (Supplementary Table S1). Cybb transcript levels were normalized to *Gapdh* and expressed relative to WT macrophages using the 2^−ΔΔCt^ method. Data represent technical duplicates.

## Acknowledgements

We thank James Imlay, Benjamin Ezraty, Laurent Aussel, Jason Yang, Severin Ronneau, and Yael Litvak for discussions. We thank Keegan Owsley and Aaron Middlebrook (BD Biosciences, now part of Waters Corporation) for advice on gating strategy design. We thank the Weizmann Institute Flow Cytometry Unit for technical assistance. We thank members of the Roi Avraham lab for helpful discussions.

## Author contributions

R.A. and C.C.M conceptualized and designed the study. C.C.M. designed and performed experiments, and analyzed data. D.S. and L.M.S. contributed the image-enabled cell sorting platform and provided technical guidance on ICS experimental design. D.S. performed image-enabled cell sorting. N.B.B.-M. performed bioinformatic analysis. H.B.-A., A.G.A., S.R.K., and S.B.-H. contributed to strain construction and infection experiments. R.A. and C.C.M. wrote the manuscript with input from all authors.

## Data availability

All sequencing data generated in this manuscript have been deposited in the Gene Expression Omnibus (GEO) under the series GSE341775 for the nTraDIS data and under the series GSE341773 for the RNA-seq data.

