## Supplementary figures for "Image-based transposon screening reveals a flavin reductase that restrains intracellular *Salmonella* replication in macrophages"

### Fig. S1

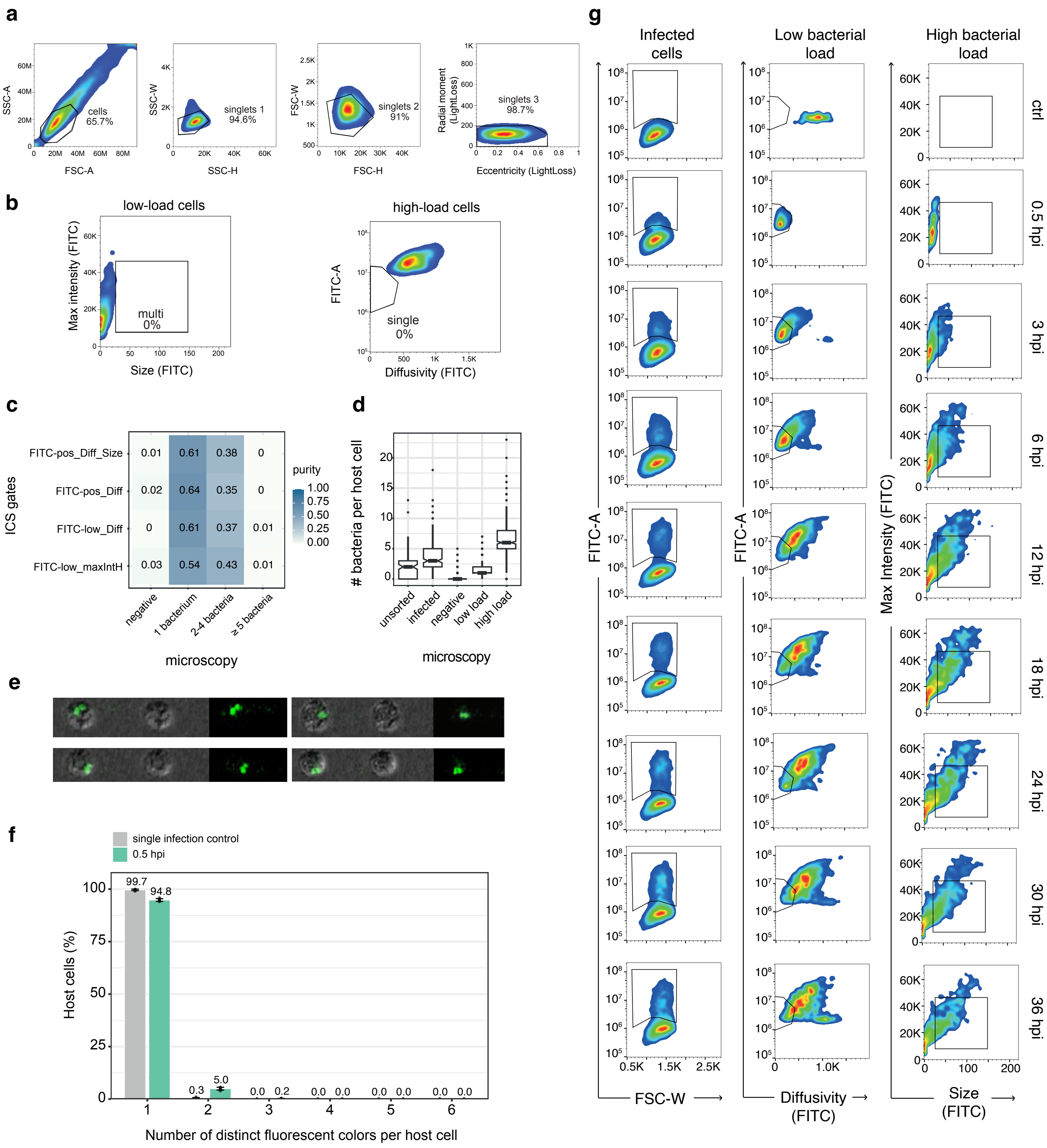
a
cells 65.7%
singlets 1 94.6%
singlets 2 91%
singlets 3 98.7%
SSC-A
FSC-A
SSC-W
SSC-H
FSC-W
FSC-H
Radial moment (LightLoss)
Eccentricity (LightLoss)
b
low-load cells
high-load cells
multi 0%
single 0%
Max intensity (FITC)
Size (FITC)
FITC-A
Diffusivity (FITC)
c
ICS gates
FITC-pos_Diff_Size 0.01 0.61 0.38 0
FITC-pos_Diff 0.02 0.64 0.35 0
FITC-low_Diff 0 0.61 0.37 0.01
FITC-low_maxIntH 0.03 0.54 0.43 0.01
negative
1 bacterium
2-4 bacteria
≥5 bacteria
microscopy
purity
d
### bacteria per host cell
unsorted
infected
negative
low load
high load
microscopy
e
f
single infection control
0.5 hpi
Host cells (%)
Number of distinct fluorescent colors per host cell
g
Infected cells
Low bacterial load
High bacterial load
ctrl
0.5 hpi
3 hpi
6 hpi
12 hpi
18 hpi
24 hpi
30 hpi
36 hpi
FITC-A
FSC-W
Diffusivity (FITC)
Max Intensity (FITC)
Size (FITC)

**Fig. S1: Related to Figure 1. ICS gating, purity controls, and infection time course. a,** Sequential gating isolates single, intact macrophages. Pre-gating sequence: cells (SSC-A/FSC-A), singlet discrimination (SSC and FSC width/height), and final singlet selection (radial moment/eccentricity). **b,** The low- and high-load gates select mutually exclusive cells. Cells within the low-load gate, projected onto the high-load axes (FITC max intensity/size, left panel), and cells within the high-load gate, projected onto the low-load axes (FITC-A/diffusivity, right panel); 0% of cells in each gate fall within the other. **c,** Comparison of candidate gate definitions identifies the best-performing low-load gate. Purity heatmap of alternative gate definitions (rows) scored against microscopy load categories (columns); the definition with the highest low-load purity was selected for sorting. 228-589 cells were scored per gate definition. **d,** Bacteria per host cell, scored by microscopy, across sorted categories (unsorted, infected, negative, low/high load). **e,** Unresolved bacterial pairs account for contamination of the low-load gate. Representative ICS images illustrating bacterial pairs aligned along the imaging axis that cannot be resolved without z-sectioning. **f,** Low-MOI infection yields predominantly single-bacterium entry. X-axis is the number of distinct fluorescent colors per macrophage, used as a proxy for independent bacterial entries, measured at 0.5 hpi (MOI 2:1) and compared to a single-infection control. **g,** Intracellular replication proceeds heterogeneously over the infection time course. Time-resolved ICS density plots for infected cells, low-load, and high-load gates from control through 0.5–36 hpi.

**Fig. S2**

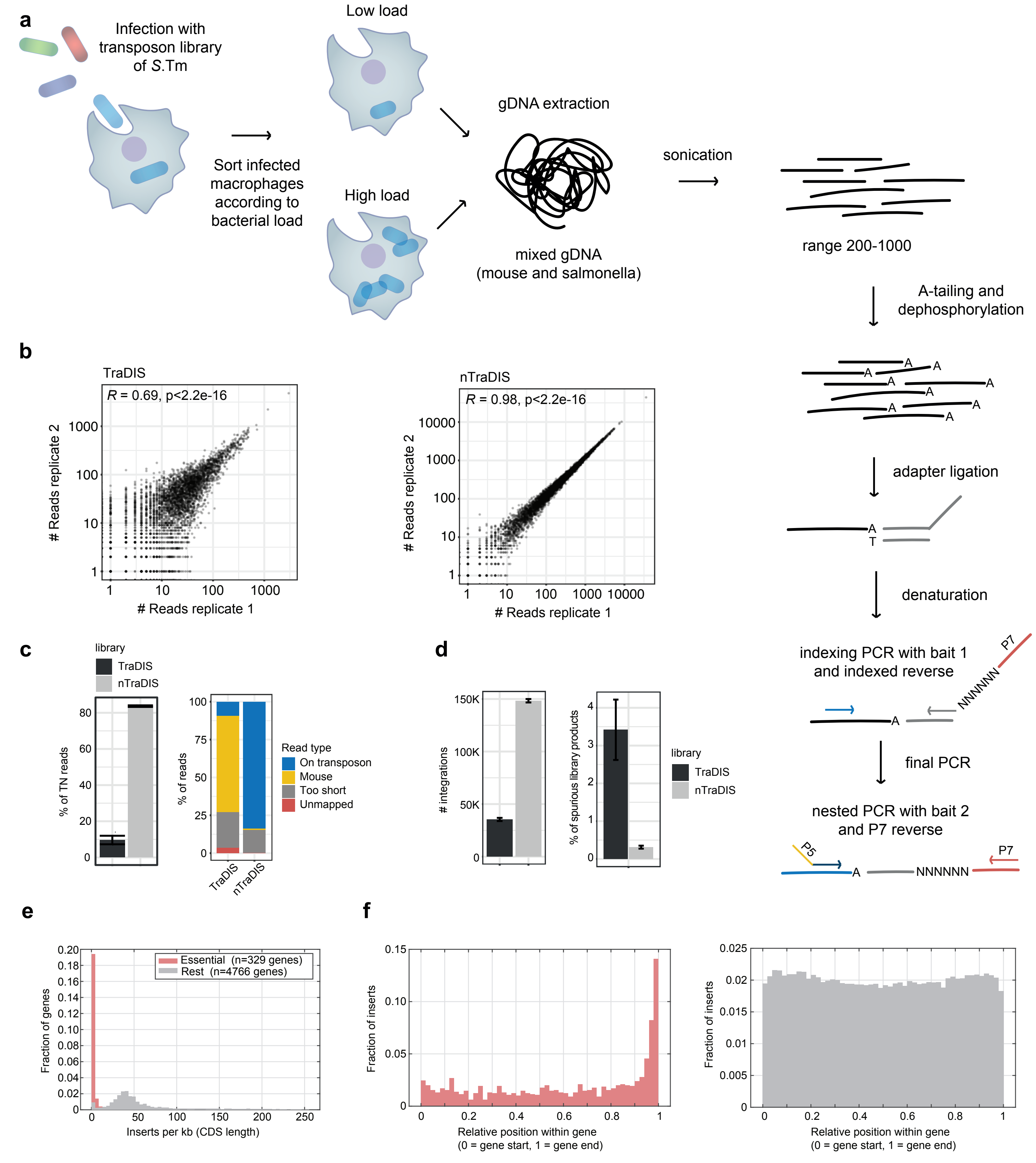

a
Infection with transposon library of S.Tm
Sort infected macrophages according to bacterial load
Low load
High load
gDNA extraction
mixed gDNA (mouse and salmonella)
sonication
range 200-1000
A-tailing and dephosphorylation
adapter ligation
denaturation
indexing PCR with bait 1 and indexed reverse
NNNNNN
P7
final PCR
nested PCR with bait 2 and P7 reverse
P5
b
TraDIS
R = 0.69, p<2.2e-16
nTraDIS
R = 0.98, p<2.2e-16
### Reads replicate 2
### Reads replicate 1
c
library
TraDIS
nTraDIS
% of TN reads
% of reads
Read type
On transposon
Mouse
Too short
Unmapped
d
### integrations
% of spurious library products
e
Essential (n=329 genes)
Rest (n=4766 genes)
Fraction of genes
Inserts per kb (CDS length)
f
Fraction of inserts
Relative position within gene
(0 = gene start, 1 = gene end)

**Fig. S2: Related to Figure 2. The nTraDIS protocol improves transposon read recovery from infected macrophages. a,** Outline of the nested TraDIS (nTraDIS) library preparation. Schematic of the workflow: sorting of infected macrophages by load, mixed (mouse + Salmonella) gDNA extraction, sonication, A-tailing/dephosphorylation, adapter ligation, indexing PCR, and nested final PCR. **b,** nTraDIS improves reproducibility between technical replicates. Correlation of insertion read counts between technical replicates for TraDIS (R = 0.69) and nTraDIS (R = 0.98). **c,** nTraDIS increases on-target transposon read recovery and reduces host contamination. On-target transposon read fraction (left) and read-type composition (on-transposon, mouse, too short, unmapped; right) for TraDIS versus nTraDIS. **d,** nTraDIS recovers more unique insertions with fewer spurious products. Unique transposon insertions recovered (left) and percentage of spurious library products (right) for the two protocols. **e**, Essential genes are depleted of insertions. Distribution of insertion density (unique insertions per kb of CDS) for genes previously annotated as essential in the literature [41] (n = 329) versus all other genes (n = 4,766, grey); essential genes cluster at near-zero insertion density. **f**, Residual insertions in essential genes localize to the 3′ end of the coding sequence. Relative position of insertions within the gene body (0 = gene start, 1 = gene end) for essential genes (left) and non-essential genes (right); insertions in essential genes are enriched toward the gene end, where they are least likely to disrupt function, whereas non-essential genes show uniform coverage.

### Fig. S3

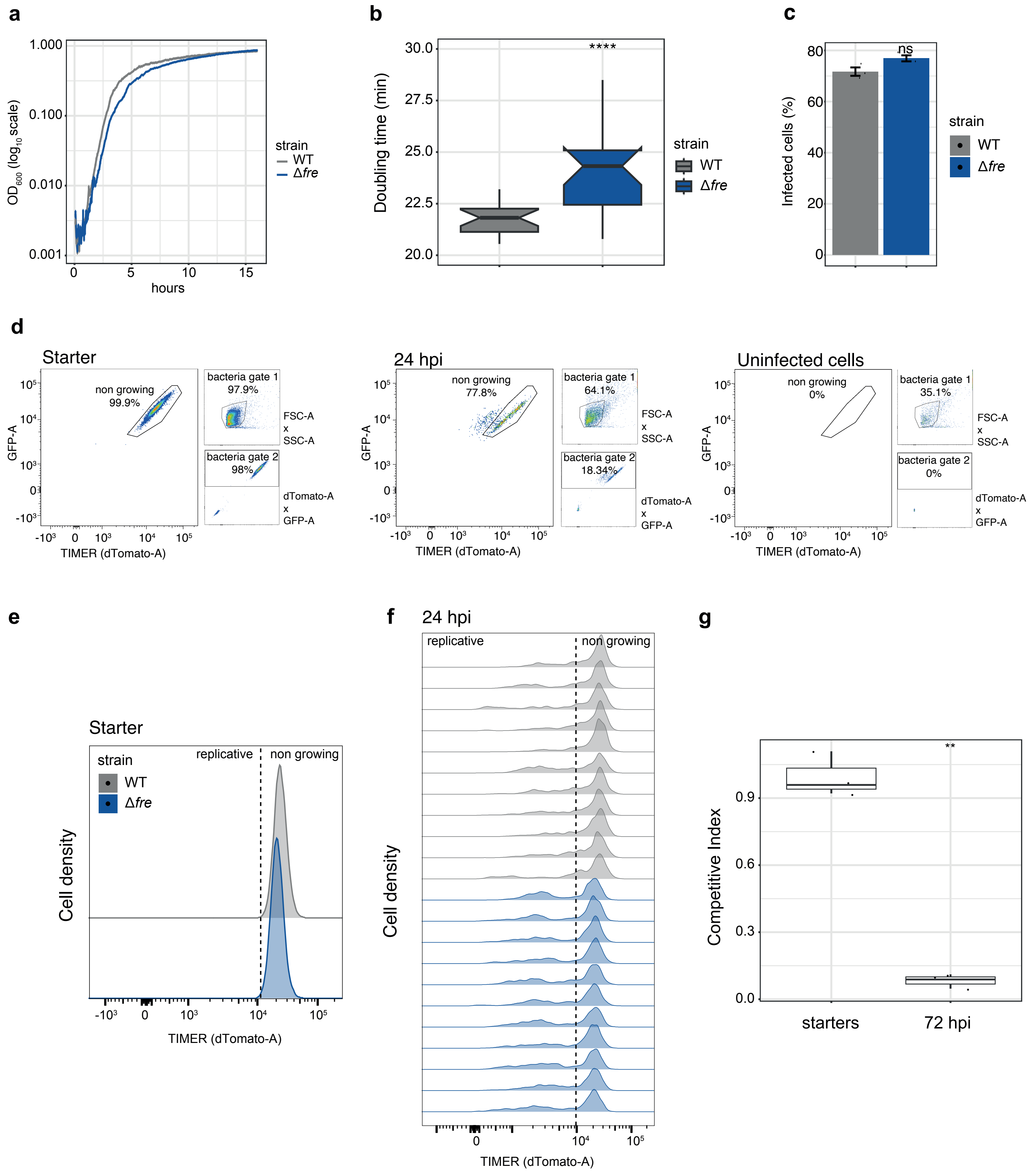

a
1.000
0.100
0.010
0.001
OD600 (log10 scale)
hours
strain
WT
Δfre
b
****
Doubling time (min)
strain
WT
Δfre
c
ns
Infected cells (%)
strain
WT
Δfre
d
Starter
non growing
99.9%
bacteria gate 1
97.9%
bacteria gate 2
98%
FSC-A x SSC-A
dTomato-A x GFP-A
GFP-A
TIMER (dTomato-A)
24 hpi
non growing
77.8%
bacteria gate 1
64.1%
bacteria gate 2
18.34%
Uninfected cells
non growing
0%
bacteria gate 1
35.1%
bacteria gate 2
0%
e
Starter
strain
WT
Δfre
replicative
non growing
Cell density
TIMER (dTomato-A)
f
24 hpi
replicative
non growing
Cell density
TIMER (dTomato-A)
g
**
Competitive Index
starters
72 hpi

**Fig. S3: Related to Figure 3. Additional characterization of the Δ*fre* replication phenotype.** **a,** Δ*fre* grows more slowly than WT in LB. Growth ($\log_{10}$ $OD_{600}$) of WT (grey) and Δ*fre* (blue) in LB broth over time. **b,** Δ*fre* has a longer doubling time than WT in LB. Doubling times for WT and Δ*fre* (n = 24; t-test: ****, P ≤ 0.0001). **c,** WT and Δ*fre* invade macrophages comparably. Percentage of infected cells for WT and Δ*fre* (mean ± SEM; n = 3; t-test: ns, P > 0.05). **d,** Gating strategy for the TIMER replication reporter. TIMER reporter gating in starter, 24 hpi, and uninfected-cell controls (GFP-A versus dTomato-A, bacteria gates). **e,** WT and Δ*fre* are equivalent in the starter inoculum. TIMER distributions of the starter inoculum (WT in grey, Δ*fre* in blue). **f,** WT and Δ*fre* reproducibly diverge by 24 hpi. Per-replicate replicative/non-growing TIMER (dTomato-A) distributions at 24 hpi. **g,** Δ*fre* is outcompeted by WT during systemic infection. Competitive index (Δ*fre* versus WT) in the spleen at the starter input and 72 hpi (n = 3 mice; t-test: **, P ≤ 0.01).

Fig. S4

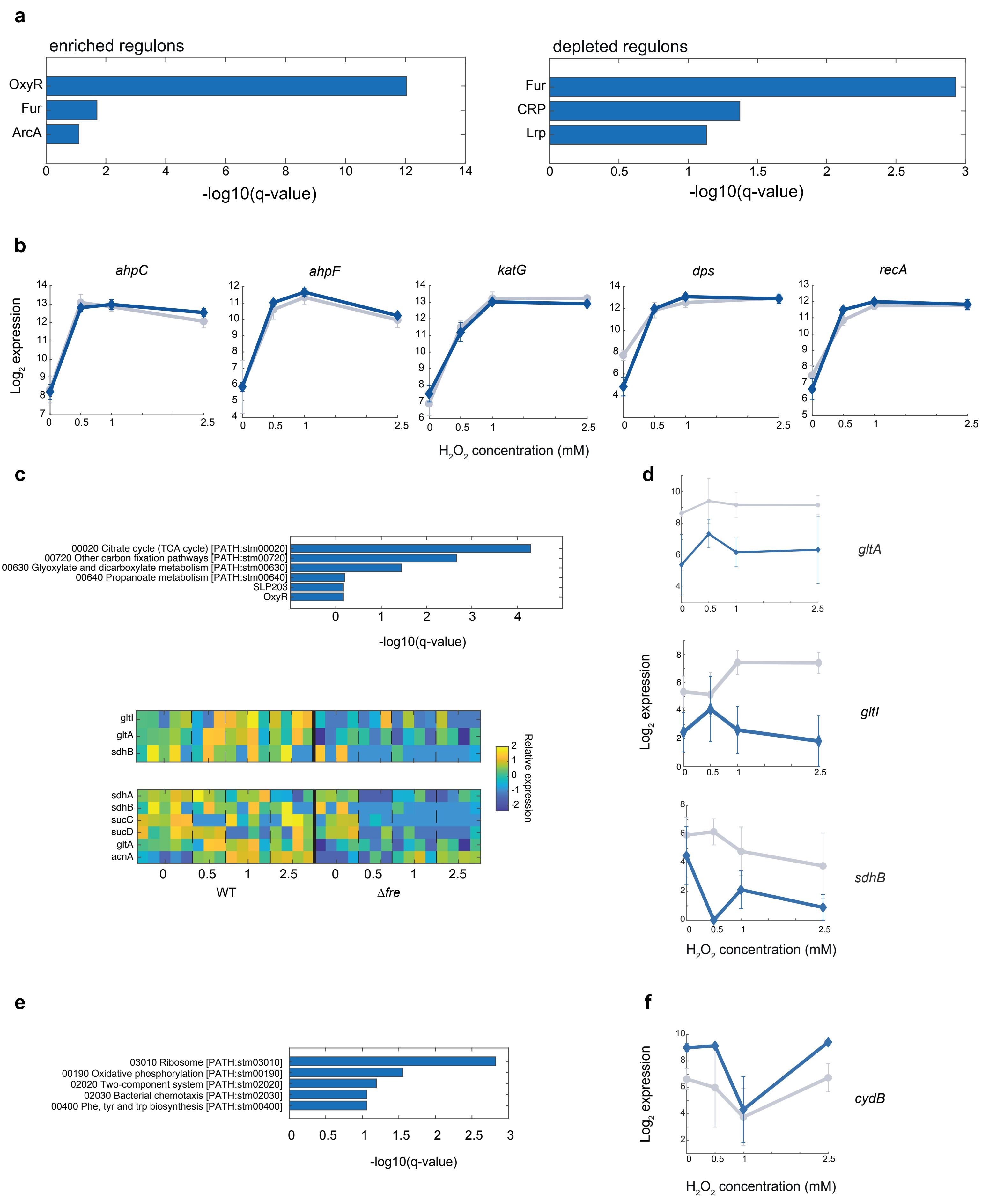
a
enriched regulons
OxyR
Fur
ArcA
-log10(q-value)
depleted regulons
Fur
CRP
Lrp
-log10(q-value)
b
ahpC
ahpF
katG
dps
recA
Log2 expression
H2O2 concentration (mM)
c
00020 Citrate cycle (TCA cycle) [PATH:stm00020]
00720 Other carbon fixation pathways [PATH:stm00720]
00630 Glyoxylate and dicarboxylate metabolism [PATH:stm00630]
00640 Propanoate metabolism [PATH:stm00640]
SLP203
OxyR
-log10(q-value)
gltI
gltA
sdhB
sdhA
sdhB
sucC
sucD
gltA
acnA
Relative expression
WT
Δfre
d
gltA
gltI
sdhB
Log2 expression
H2O2 concentration (mM)
e
03010 Ribosome [PATH:stm03010]
00190 Oxidative phosphorylation [PATH:stm00190]
02020 Two-component system [PATH:stm02020]
02030 Bacterial chemotaxis [PATH:stm02030]
00400 Phe, tyr and trp biosynthesis [PATH:stm00400]
-log10(q-value)
f
cydB
Log2 expression
H2O2 concentration (mM)

**Fig. S4: Related to Figure 4. Transcriptional response to $H_2O_2$ in WT and Δ*fre*. a,** The peroxide response is dominated by OxyR and Fur regulons in both genotypes. Regulons significantly enriched (left) and depleted (right) in the $H_2O_2$ response. **b,** The direct $H_2O_2$-sensing core of the OxyR regulon is induced normally in Δ*fre*. Dose-response of canonical OxyR targets (*ahpC*, *ahpF*, *katG*, *dps*, *recA*) in WT versus Δ*fre*; mean ± SD. **c,** Genes repressed in Δ*fre* are enriched for the TCA cycle. Top, KEGG pathway enrichment among genes repressed in Δ*fre*. Bottom, heatmaps of relative expression across the $H_2O_2$ dose series in WT and Δ*fre* for genes annotated to the OxyR regulon (*gltI*, *gltA*, *sdhB*; top) and to the TCA cycle (*sdhAB*, *sucCD*, *gltA*, *acnA*; bottom). **d,** Repression in Δ*fre* is consistent across the dose series. Per-gene dose-response traces for *gltI*, *gltA*, and *sdhB* in WT versus Δ*fre*; mean ± SD. **e,** Genes induced in Δ*fre* are enriched for ribosome, oxidative phosphorylation, two-component system, chemotaxis, and aa biosynthesis. KEGG pathway enrichment among genes induced in Δ*fre*. **f,** *cydB* is constitutively elevated in Δ*fre*. Transcript levels ($log_2$ expression) of *cydB* as a function of $H_2O_2$ concentration in WT (grey) and Δ*fre* (blue); mean ± SD.

**Fig. S5**

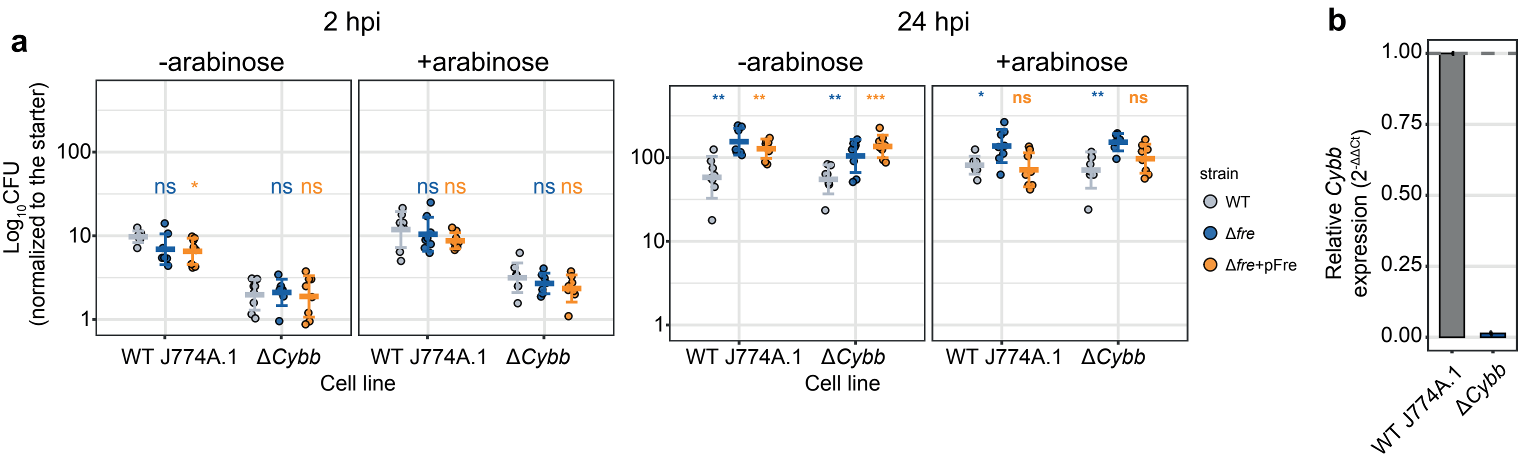

a
2 hpi
24 hpi
-arabinose
+arabinose
Log$_{10}$CFU (normalized to the starter)
100
10
1
ns
*
ns ns
WT J774A.1
ΔCybb
Cell line
**
***
strain
WT
Δfre
Δfre+pFre
b
Relative Cybb expression ($2^{-\Delta\Delta Ct}$)
1.00
0.75
0.50
0.25
0.00

**Fig. S5: Related to Figure 5. The Δ*fre* replication advantage does not require host NADPH oxidase. a,** Δ*fre* out-replicates WT to a comparable extent in WT and Δ*Cybb* macrophages, and inducible complementation restores WT-like replication in both. Intracellular CFU in WT J774A.1 and Δ*Cybb* macrophages (−/+ arabinose, 2 and 24 hpi) for WT, Δ*fre*, and Δ*fre* + pFre (n = 8-16; t-test significance markers colored by strain). **b,** *Cybb* expression is lost in Δ*Cybb* macrophages. Relative *Cybb* expression ($2^{-\Delta\Delta Ct}$) in WT J774A.1 versus Δ*Cybb* (n = 3 technical replicates).
